# A thalamic reticular circuit for head direction cell tuning and spatial navigation

**DOI:** 10.1101/804575

**Authors:** Gil Vantomme, Zita Rovó, Romain Cardis, Elidie Béard, Georgia Katsioudi, Angelo Guadagno, Virginie Perrenoud, Laura MJ Fernandez, Anita Lüthi

## Abstract

To navigate in space, an animal must refer to sensory cues to orient and move. Circuit and synaptic mechanisms that integrate cues with internal head-direction (HD) signals remain, however, unclear. We identify an excitatory synaptic projection from the presubiculum (PreS) and the multisensory-associative retrosplenial cortex (RSC) to the anterodorsal thalamic reticular nucleus (TRN), so far classically implied in gating sensory information flow. *In vitro*, projections to TRN involved AMPA/NMDA-type glutamate receptors that initiated TRN cell burst discharge and feedforward inhibition of anterior thalamic nuclei. *In vivo*, chemogenetic anterodorsal TRN inhibition modulated PreS/RSC-induced anterior thalamic firing dynamics, broadened the tuning of thalamic HD cells, and led to preferential use of allo-over egocentric search strategies in the Morris water maze. TRN-dependent thalamic inhibition is thus an integral part of limbic navigational circuits wherein it coordinates external sensory and internal HD signals to regulate the choice of search strategies during spatial navigation.

## Introduction

To reach a goal, search for food, or avoid a predator, navigation in space is essential. To do so accurately, animals rely on sensory landmarks in the environment to monitor and adapt their path and body orientation in space. Sensory cortices elaborate information they receive from the sensory organs, and they interact with widespread thalamo-cortico-hippocampal networks that contain internal representations of space and body orientation to guide navigation (Hinman et al., 2018; Vélez-Fort et al., 2018). In the interest of survival in a constantly changing world, salient environmental cues have to be integrated rapidly with spatial signals so that navigational strategies can be adapted prior to full cortical elaboration. Accordingly, important integrative steps enabling sensory guided spatial navigation may be engaged at subcortical levels (Hinman et al., 2018; Knudsen, 2018). A major site for subcortical gating of external sensory stimuli is the inhibitory thalamic reticular nucleus (TRN) that shows a unique anatomical positioning at the interface between sensory thalamic nuclei and cortex (Crabtree, 2018; Pinault, 2004; Scheibel and Scheibel, 1966). Activity in TRN controls the gain of sensory inputs (Le Masson et al., 2002), sharpens receptive fields in thalamic sensory nuclei (Lee et al., 1994; Soto-Sánchez et al., 2017), underlies sensory selection in divided attentional tasks (Ahrens et al., 2015; Wimmer et al., 2015), and is involved in sensory induced flight responses (Dong et al., 2019).

To date, the TRN has not been implied in the gating of spatial information. Lesion studies, however, suggest that TRN contributes to covertly directing a rat’s self-orientation to the target stimulus, such that orienting movements can be rapidly executed (Weese et al., 1999). Moreover, anterior thalamic nuclei (ATN) are part of the brain’s navigational system (Dumont and Taube, 2015), and there is anatomical evidence in rodent that anterodorsal TRN innervates ATN (Gonzalo-Ruiz and Lieberman, 1995a, b; Lozsádi, 1995; Pinault and Deschênes, 1998; Scheibel and Scheibel, 1966), although this has been questioned in cat (Paré et al., 1987). The anterodorsal (AD) thalamic nucleus, part of the ATN, contains a large proportion of HD cells tuned to the direction of the rodent’s head in space (Taube, 1995). The HD signal is generated within dorsal tegmental and lateral mammillary nuclei based primarily on vestibular signals and is then relayed to AD and from there to PreS (Sharp et al., 2001a; Sharp et al., 2001b; Stackman and Taube, 1998). Although the TRN has been proposed to be part of HD circuits (Peyrache et al., 2019), the underlying functional anatomy remains elusive. Possible equivalences and differences to the canonical sensory TRN-thalamocortical circuits thus remain speculative. Here, we hypothesized that if the TRN is to mediate subcortical sensory gating effectively, it should serve as an entry point for information flow to ATN to control the processing of HD signals.

The anterior thalamic HD representation is updated rapidly, within < 100 ms by external visual landmarks (Zugaro et al., 2003), to which direct input from the dorsal presubiculum (dPreS) (Goodridge and Taube, 1997), indirect input from dPreS via the lateral mammillary nucleus (Yoder et al., 2015) and the retrosplenial cortex (RSC) (Clark et al., 2010) are thought to contribute. Both PreS and RSC are reciprocally connected (van Groen and Wyss, 1990) and receive afferents from ATN, primary and secondary visual cortex, integrating information relevant for egocentric and allocentric, external cue-guided, navigation (Clark et al., 2018; Dumont and Taube, 2015; Mitchell et al., 2018; Simonnet and Fricker, 2018). Behaviorally, lesions of dPreS compromise rapid orienting behaviors based on landmarks (Yoder et al., 2019), whereas RSC lesions lead to multiple deficits in spatial navigation and memory formation (Clark et al., 2018; Mitchell et al., 2018). Although there is evidence for a topographically organized cortical feedback from RSC to rat and monkey anterodorsal TRN (Cornwall et al., 1990; Lozsádi, 1994; Zikopoulos and Barbas, 2007), the nature of this corticothalamic communication has never been characterized. Indeed, current models of HD circuits involving ATN, dPreS and RSC (Dumont and Taube, 2015; Perry and Mitchell, 2019; Peyrache et al., 2017; Simonnet and Fricker, 2018) and of the brain’s ‘limbic’ navigational system (Bubb et al., 2017) largely disregard a functionally integrated TRN. In spite of this gap of knowledge, the notion of a ‘limbic’ anterior TRN has been proposed recently (Halassa et al., 2014; Zikopoulos and Barbas, 2012).

In this study, we combined tracing techniques, *in vitro* and *in vivo* electrophysiological recordings together with a spatial navigation task to probe the synaptic integration and the function of TRN in the communication between PreS, RSC and ATN.

## Results

### RSC and PreS send topographically organized projections to ATN and TRN

To determine afferent projections to the anterodorsal portion of the TRN, we injected small volumes (50-100 nl) of red retrobeads into anterodorsal TRN of C57BL6/J mice (4-8-week-old) and identified sites of red punctate fluorescent labeling 5 – 7 days later. Four out of 19 injections were restricted to the dorsal portion of the anterior TRN, as verified by parvalbumin (PV)- immunostaining (Fig. 1A1, Suppl. Fig. 1). The size of injection sites was comparable to previous reports using nanobead injections in TRN (Antal et al., 2014). We inspected brain areas in which punctate retrobead labeling was present and clearly separated from the bulk of beads around the injection site. We found punctate staining in the adjacent AD, laterodorsal (LD) and in the centrolateral (CL) nuclei of the thalamus (Fig. 1A2, identified in 3 out of 4 injections), consistent with prior tracing studies (Gonzalo-Ruiz and Lieberman, 1995a, b; Lozsádi, 1995; Pinault and Deschênes, 1998). Labeling was also found in deep layers of prelimbic cortex that extended into infralimbic and cingulate, and, in two cases, into motor cortical areas (not shown), consistent again with a previous study in rat (Lozsádi, 1994). Our attention was drawn to a distinct stretch of puncta extending from parahippocampal regions into RSC that was present for all four injections (Fig. 1A2, Suppl. Fig. 1). Labeling included in particular the deep layers of the PreS that is interposed between the subiculum, the parasubiculum and the RSC (Ding, 2013; Simonnet and Fricker, 2018).

**Figure 1.**
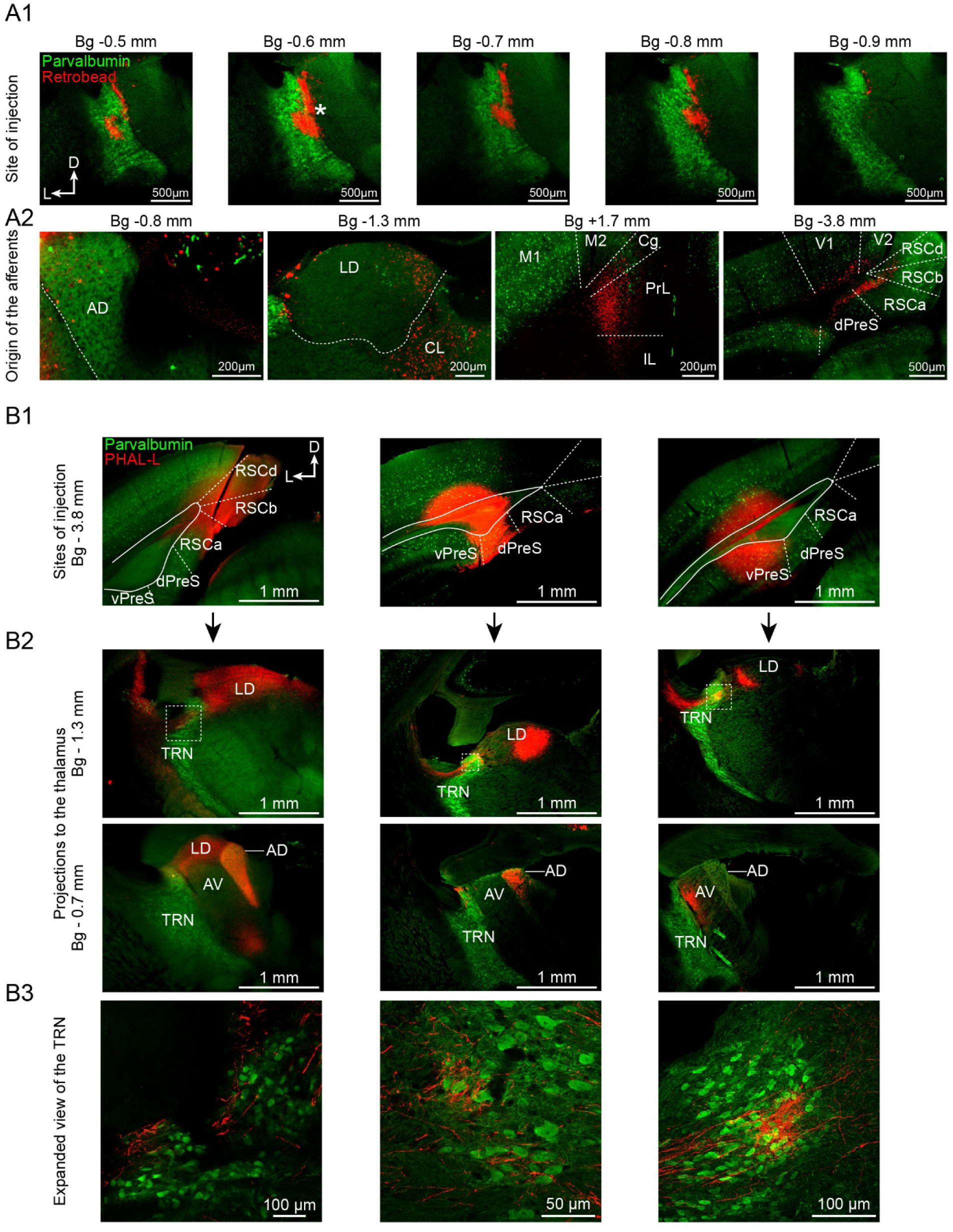
The RSC and the PreS send topographically organized projections to ATN and TRN. **(A1)** Epifluorescent micrographs of mouse coronal brain sections showing a retrobead (red) injection site (*) into the anterior portion of the TRN, which spread ∼300 μm along the anteroposterior extent of the TRN (immunostained for PV, green). Bg, Bregma. **(A2)** Epifluorescent micrographs showing retrogradely labeled brain regions. Anterodorsal thalamus (AD) – Laterodorsal thalamus (LD) – Centrolateral thalamus (CL) – Cingulate cortex (Cg) – Prelimbic/Infralimbic cortex (PreL/IL) – dorsal Presubiculum (dPreS) – Retrosplenial cortex (RSC) – Visual cortex (V1/V2) – Motor cortex (M1/M2). **(B1)** Epifluorescent micrographs of 3 different injection sites of PHAL-L (red) into (from left to right) RSC, dPreS and ventral PreS (vPreS). Green, PV+ neurons. **(B2)** Epifluorescent micrographs of coronal sections in ATN at Bg -1.3 mm (top) and -0.7 mm (bottom). Note labeled fibers visible in the anterodorsal TRN (dotted squares). **(B3)** Expanded confocal microscopy views of areas indicated by dotted squares in B2, AV, anteroventral thalamus.

We next used the anterograde tracer, *Phaseolus vulgaris*-leucoagglutinin (PHAL-L), to further assess anatomical projections specifically from RSC and PreS to anterior TRN. Through a panel of injections (n = 20) that targeted restricted portions of RSC, dPreS and ventral PreS (vPreS) (Fig. 1B1), we noted a nucleus-specific labelling pattern in the LD, AD, and anteroventral (AV) thalamus (the later is also part of ATN) (Fig. 1B2). Injections centered within the RSC labeled large portions of AD and LD, while sparing AV, whereas PreS-centered injections covered more restricted portions of LD, AD and AV. vPreS injections labeled the most lateral portion of LD and AV. Using confocal microscopy to follow the path of PreS/RSC fibers on their way to ATN and LD (together referred as ATN+), we realized that these traversed the TRN as a bundle that showed marked arborizations within the most anterodorsal portion of TRN (Fig. 1B3). Arborizations surrounded the TRN somata in dense plexuses and ran along portions of dendrites, suggesting that they formed synaptic contacts.

### The PreS/RSC establishes functional excitatory synapses onto TRN

To quantify these anatomical observations in terms of possible functional synapses, we used whole-cell patch-clamp recordings in acute coronal slices from brains of mice injected with AAV1-CaMKIIa-ChR2-EYFP into PreS/RSC 3 – 5 weeks earlier (Fig. 2A). Cells patched within anterodorsal TRN showed rebound burst behavior, as recognizable by repetitive high-frequency bursts of action potentials after brief hyperpolarization, similar to posterior sensory TRN cells (Fig. 2B) (Fernandez et al., 2018). Electrical properties were also comparable to those of their posterior counterparts that are innervated by the primary somatosensory cortex S1 (Fernandez et al., 2018; Vantomme et al., 2019), but cells produced less repetitive bursts (Fig. 2C). Cells in AD, AV and LD showed properties typical for dorsal thalamocortical neurons, notably the presence of only a single rebound burst discharge (Huguenard, 1996) (Suppl. Fig. 2).

**Figure 2.**
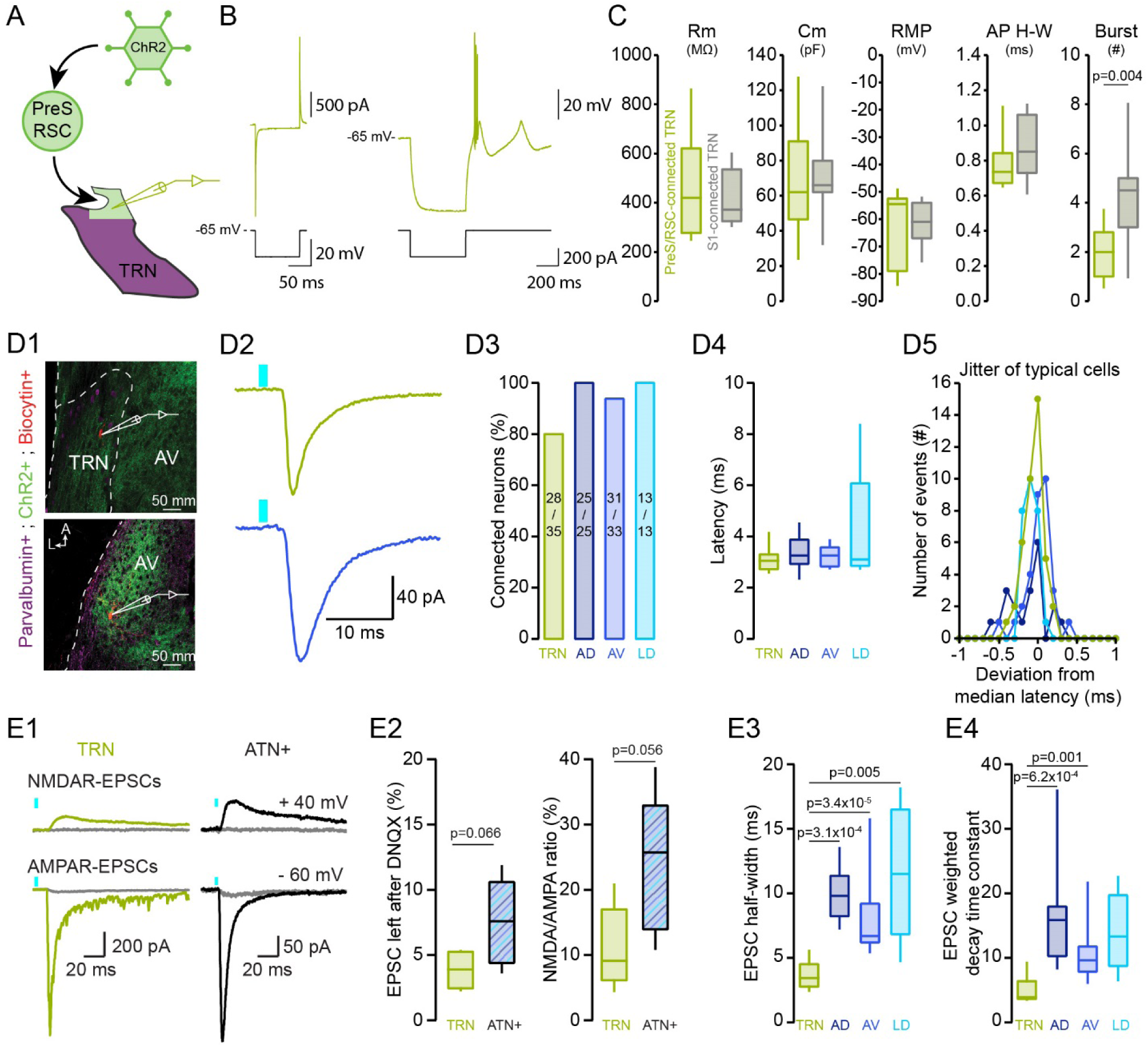
The PreS/RSC establishes functional excitatory synapses onto TRN. **(A)** Scheme of viral injections (AAV1-CamKIIa-ChR2-EYFP) into PreS/RSC followed by whole-cell patch-clamp recordings. **(B)** Responses of a PreS/RSC-connected TRN neuron (green) to a hyperpolarizing step in voltage-clamp (Left) and to negative current injection in current-clamp (Right). **(C)** Box-and-whisker plots of cellular properties of PreS/RSC-connected (green, n = 16) and, for comparison, S1-connected TRN neurons (grey, n = 11). Data from S1-connected TRN neurons re-used from a previous study (Fernandez et al., 2018). From left to right: membrane resistance (Rm), membrane capacitance (Cm), resting membrane potential (RMP), action potential (AP) half-width (H-W), burst number. Mann-Whitney U tests were used for comparing Rm, RMP and AP H-W, Student’s *t* tests for Cm and Burst number. **(D1)** Confocal micrographs of 300 μm-thick mouse brain sections showing the whole-cell recorded TRN (top) and AV (bottom) neurons filled with neurobiotin (red). Green, ChR2-EYFP-expressing PreS/RSC afferents, magenta, PV+ TRN cells. **(D2)** Current responses of TRN (top) and AV (bottom) neurons to optogenetic activation (blue bars, 1 ms, 3.5 mW power, 455 nm) of PreS/RSC afferents, recorded at -60 mV. **(D3)** Connectivity histogram, calculated as the fraction (in %) of neurons responding to optogenetic stimulation. **(D4)** Box-and-whisker plot of response latencies (calculated from LED onset to response onset) in the TRN (n = 12), AD (n = 16), AV (n = 16) and LD (n = 6). Mann-Whitney U tests and Bonferroni correction: α = 0.0083. **(D5)** Jitter of response latencies (plotted as the deviation from mean for all individual response events) in one cell from TRN, AD, AV and LD across all stimulation trials. **(E1)** Pharmacological analysis of typical evoked excitatory postsynaptic currents (EPSCs) in TRN and ATN+, showing AMPA- and NMDA-EPSCs and their suppression by DNQX (40 μM) and APV (100 μM), respectively (superimposed grey traces). **(E2)** Box-and-whisker plots of DNQX effects (left, in % of original response amplitude, n = 5 for both TRN and ATN+) and of NMDA/AMPA ratios (right). Values of p from Student’s *t* tests. **(E3)** Box-and-whisker plot of EPSC half-widths for TRN (n = 7), AD (n = 8), AV (n = 14) and LD (n = 5). Mann-Whitney U tests and Bonferroni correction: α = 0.0083. Statistically significant p values are indicated. **(E4)** Box-and-whisker plot of the EPSC weighted decay time constant in TRN (n = 7), AD (n = 8), AV (n = 14) and LD (n = 5). Same statistical analysis as E3.

Optogenetic stimulation of PreS/RSC fibers was applied while recording from voltage-clamped neurons of the anterodorsal TRN and of AD, AV and LD (Fig. 2D,E). The location of cells within the different thalamic nuclei was evident while guiding the patch pipette to the target region and was confirmed in a subgroup of cells through perfusion with neurobiotin and *post-hoc* recovery (n = 33/106) (Fig. 2D1). Rapid synaptic inward currents were elicited in > 80 % of all recorded cells for all areas (Fig. 2D2,D3). Synaptic currents were time-locked to the stimulus, with a fixed and short latency to response onset and sub-millisecond jitter (Fig. 2D4,D5). Response latency was inversely proportional to light intensity (Suppl. Fig. 2), which is consistent with an action potential-dependent mode of synaptic transmission (Gjoni et al., 2018). There is thus a direct, monosynaptic connection from PreS/RSC to anterodorsal TRN and to AD, AV and LD.

Light-evoked postsynaptic currents (EPSCs) were mediated by glutamatergic synaptic receptors, as verified in a subset of 5 TRN and 5 neurons of AD, AV or LD (jointly referred to here as ATN+) (Fig. 2E1). Thus, the AMPA receptor antagonist 6,7-Dinitroquinoxaline-2,3(1H,4H)-dione (DNQX, 40 μM, bath-application) reduced responses by > 90 % at -60 mV (Fig. 2E1,E2). The small remaining current component was abolished by the NMDA receptor antagonist DL-2-Amino-5-phosphonovaleric acid (APV, 100 µM, bath-application), as measured at +40 mV (Fig. 2E1, E2). NMDA/AMPA ratios were comparable to previous studies on corticoreticular and corticothalamic synapses in the sensory sectors of TRN (Fernandez et al., 2017). Moreover, the TRN-EPSCs had a twice-shorter half-width than ATN+-EPSCs (Fig. 2E3) and a faster decay time (Fig. 2E4). PreS/RSC inputs thus convey a phasic excitatory input onto anterodorsal TRN cells.

### PreS/RSC establishes strong unitary connections with driver characteristics onto anterodorsal TRN

TRN and ATN+ neurons were robustly innervated by PreS/RSC afferents, with light-evoked EPSC amplitudes ranging from -25 pA to -1157 pA at high light intensities, although there were nucleus-specific differences (Fig. 3A). Both large and small EPSCs were obtained in slices from the same animals, excluding variable viral transduction as a major reason for this variability. To assess how variability was based on strength and connectivity of PreS/RSC afferents, we used minimal optogenetic stimulation through reducing light intensity to variably evoke failures and successful responses at comparable rates (mean failure rate 47±3 %, at 0.28±0.05 mW) (Fig. 3B1). Unitary PreS/RSC EPSCs of TRN cells were 4-to 5-fold larger than the ones established onto AD and AV cells (Fig. 3B2). Dividing the maximally evoked EPSC amplitude by the unitary one, we calculated ranges of 1 – 9 fibers for TRN, 2 – 32 fibers for AD and 8 – 86 fibers for AV. Therefore, although variable, TRN cells are, on average, targeted by a comparatively small number of fibers, but each with greater unitary strength. A large unitary response size has also been described for cortical projections onto sensory TRN (Cruikshank et al., 2010; Gentet and Ulrich, 2004; Golshani et al., 2001). To determine how many of these fibers were necessary to bring TRN cells to threshold for action potential firing, we performed cell-attached patch-clamp recording to preserve cellular integrity during PreS/RSC synaptic stimulation (Fig. 3C1). Action current numbers showed a steep sigmoidal light dependence with half-maximal values reached at 0.8 mW (Fig. 3C2,C3). Single spikes could be detected at a light intensity corresponding to the one used for minimal stimulation (0.28±0.05 mW). Subsequent whole-cell mode recording in 5 out of 6 cells confirmed that these were bursts of action potentials riding on a low-threshold calcium spike, which showed similar light dependence (half-maximal number of action potentials at 1.1 mW) (Fig. 3C1,C3).

**Figure 3.**
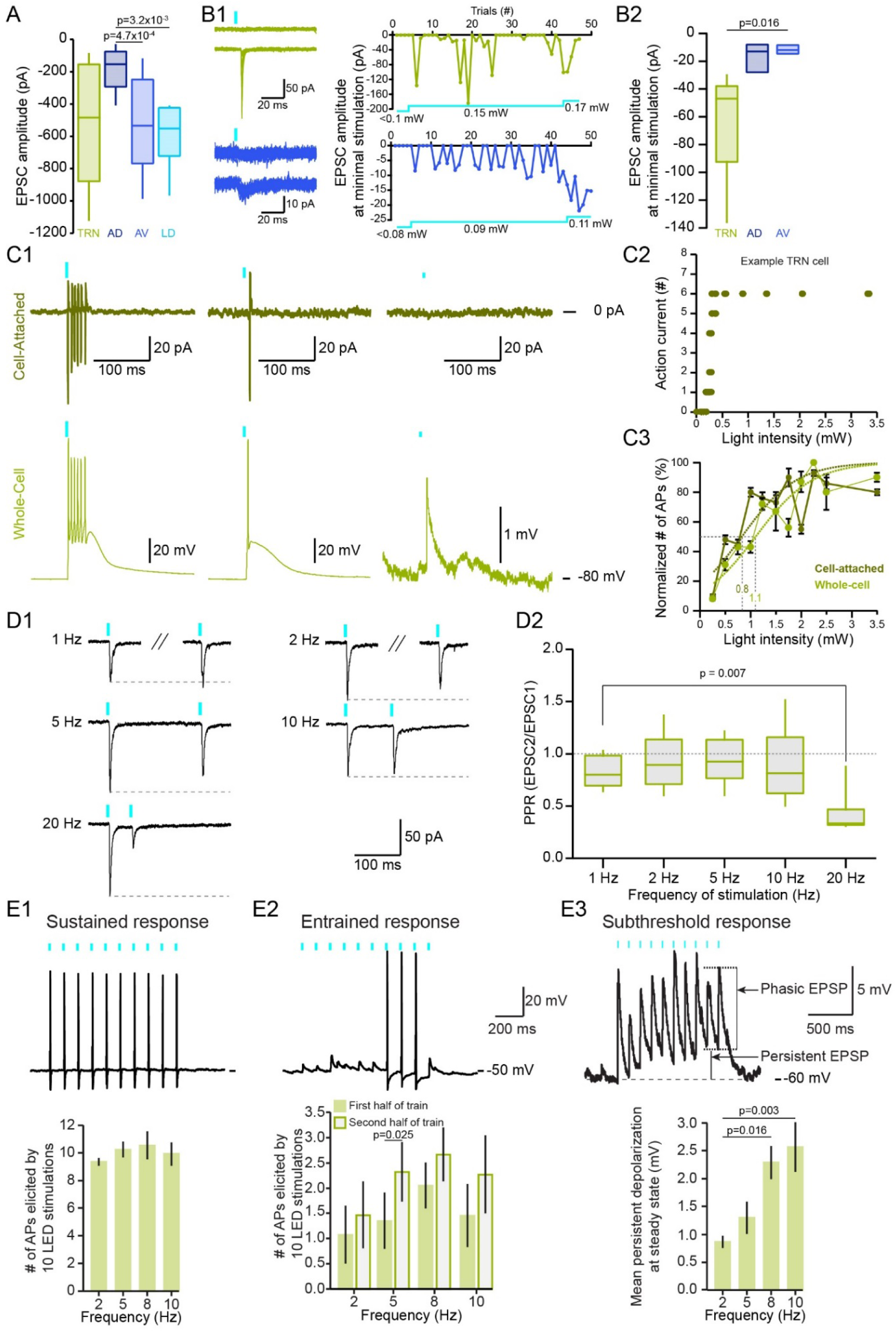
PreS/RSC establishes strong unitary connections with driver characteristics onto anterodorsal TRN. **(A)** Box-and-whisker plot of maximally evoked compound EPSC amplitudes in TRN (n = 12), AD (n = 16), AV (n = 16), LD (n = 6). The intensity of the LED was reduced to ∼20 % of the maximum in 4/16 AV and 4/6 LD cells to prevent escape currents. 1-factor ANOVA, p = 1.25×10^−3^, post hoc Student’s *t* tests with Bonferroni correction: α=0.0083. **(B1)** Minimal stimulation experiment. Left, overlay of successes and failures for a TRN and an AV neuron in one experiment. Right: Time course of the same experiment. Blue trace: intensity of the light stimulation. At minimal stimulation (0.15 mW for the TRN cell and 0.09 mW for the AV cell), the failure rate was ∼50 % (22/39 failures for the TRN neuron, 20/39 failures for the AV neuron). Increasing the light intensity brought the failure rate to 0 % (right part of the graph). **(B2)** Box-and-whisker plot of the amplitude of successfully evoked unitary EPSCs in TRN (n = 5), AD (n = 3) and AV (n = 4). Repeated Mann-Whitney U tests with Bonferroni correction: α = 0.017. **(C1)** Top: representative responses of a cell-attached TRN neuron recording exposed to maximal (left), intermediate (middle) and low (right) light intensities. Bottom: Same experiment in whole-cell current-clamp mode. **(C2)** Graph of action current number for the TRN neuron shown in C1. **(C3)** Same as in C2 for the average of all TRN neurons (cell-attached n = 6, whole-cell n = 5). Data were binned in 0.25 mW light steps. Action current number normalized to the maximum evoked in each neuron. For sigmoidal fits, see Methods. Half-maximal values are indicated. **(D1)** Representative TRN EPSCs at -60 mV upon paired-pulse stimulation at 1, 2, 5, 10 and 20 Hz. Grey dotted lines: amplitude of the first EPSC. **(D2)** Box-and-whisker plot of paired-pulse ratios (TRN: n = 16). Paired Student’s t tests or Wilcoxon signed rank-test and Bonferroni correction: α = 0.013. **(E1)** Top: typical membrane voltage response of a TRN neuron to a 10 Hz-light stimulation train. Bottom: Histogram of means (n = 7). Wilcoxon signed rank-tests and Bonferroni correction: α = 0.017. **(E2)** Top: same as in E1 for neurons responding with a subthreshold response at train onset. Bottom: Histogram of means (n = 6). Wilcoxon signed rank-test (at 2 Hz) and Paired Student’s *t* tests (at 5, 8, 10 Hz). **(E3)** Top: same as in E1 for subthreshold responses in a TRN neuron held at -60 mV. Bottom: Histogram of the mean persistent depolarization (n = 5). The persistent depolarization measured on the last 3 stimulations. 1-factor RM ANOVA, p = 0.033, post hoc paired Student’s *t* tests and Bonferroni correction: α = 0.017.

Single or few active synaptic inputs from PreS/RSC appear thus sufficient to bring TRN cells to threshold through reliable EPSP-low threshold burst coupling.

Excitatory afferents into thalamus have been divided into 2 major groups, drivers and modulators (Sherman, 2017). To start addressing the profile of PreS/RSC afferents, we determined paired-pulse ratios (PPRs) of TRN-and ATN+-EPSCs. Under our ionic conditions, PPRs remained close to 1 until at least 10 Hz (Fig. 3D1,D2, Suppl. Fig. 3). When plotting results from individual experiments, all data points clustered around 1 for 1-10 Hz. This short-term plasticity profile is markedly different from the modulatory profile of cortical input onto sensory TRN, which shows a strong paired-pulse facilitation (PPF) (Fernandez et al., 2018).

At depolarized potentials (−50 mV), where tonic discharge is prevalent, PreS/RSC afferents reliably sustained TRN discharge during stimulation trains (Fig. 3E1). Furthermore, initially subthreshold responses could become suprathreshold in the course of a train (Fig. 3E2), most likely due to temporal summation that gave rise to a persistent depolarization on top of the phasic events (Fig. 3E3). Similar results were found at PreS/RSC-ATN+ synapses (Suppl. Fig. 3).

### PreS/RSC afferents mediate feedforward inhibition onto ATN+ through recruiting burst discharge in PV- and somatostatin (Sst)-expressing TRN cells

How does TRN recruitment by PreS/RSC afferents regulate ATN+ activity? We first tested *in vitro* for Pres/RSC-triggered feedforward inhibition onto ATN+ (Fig. 4A). ATN+ cells were held at voltages to separately monitor EPSC and IPSC components (−60 mV and +15 mV) (see Methods for further details). Out of 22 ATN+ neurons innervated by PreS/RSC, 19 (9 AD, 5 AV, 5 LD) presented with a strong outward current at +15 mV, consistent with an evoked inhibitory postsynaptic current (IPSC) (Fig. 4B). The IPSC latency was higher than the EPSC latency (Fig. 4C), consistent with a disynaptic feedforward inhibition. IPSCs were mediated through GABA_A_ receptors (Fig. 4D1,D2). To demonstrate that these IPSCs were indeed mediated by anterodorsal TRN, we combined opto- and chemogenetics in VGAT-Ires-Cre mice expressing the inhibitory Designer Receptor Exclusively Activated by Designer Drugs (inhibitory DREADD, abbreviated as hM4D from here onwards) specifically in the GABAergic cells of anterodorsal TRN and ChR2 in PreS/RSC. Chemogenetic silencing of anterodorsal TRN through bath-application of the hM4D ligand clozapine N-oxide (CNO) while optogenetically activating PreS/RSC afferents indeed reduced the amplitude of the evoked IPSC (Fig. 4E1,E2).

**Figure 4.**
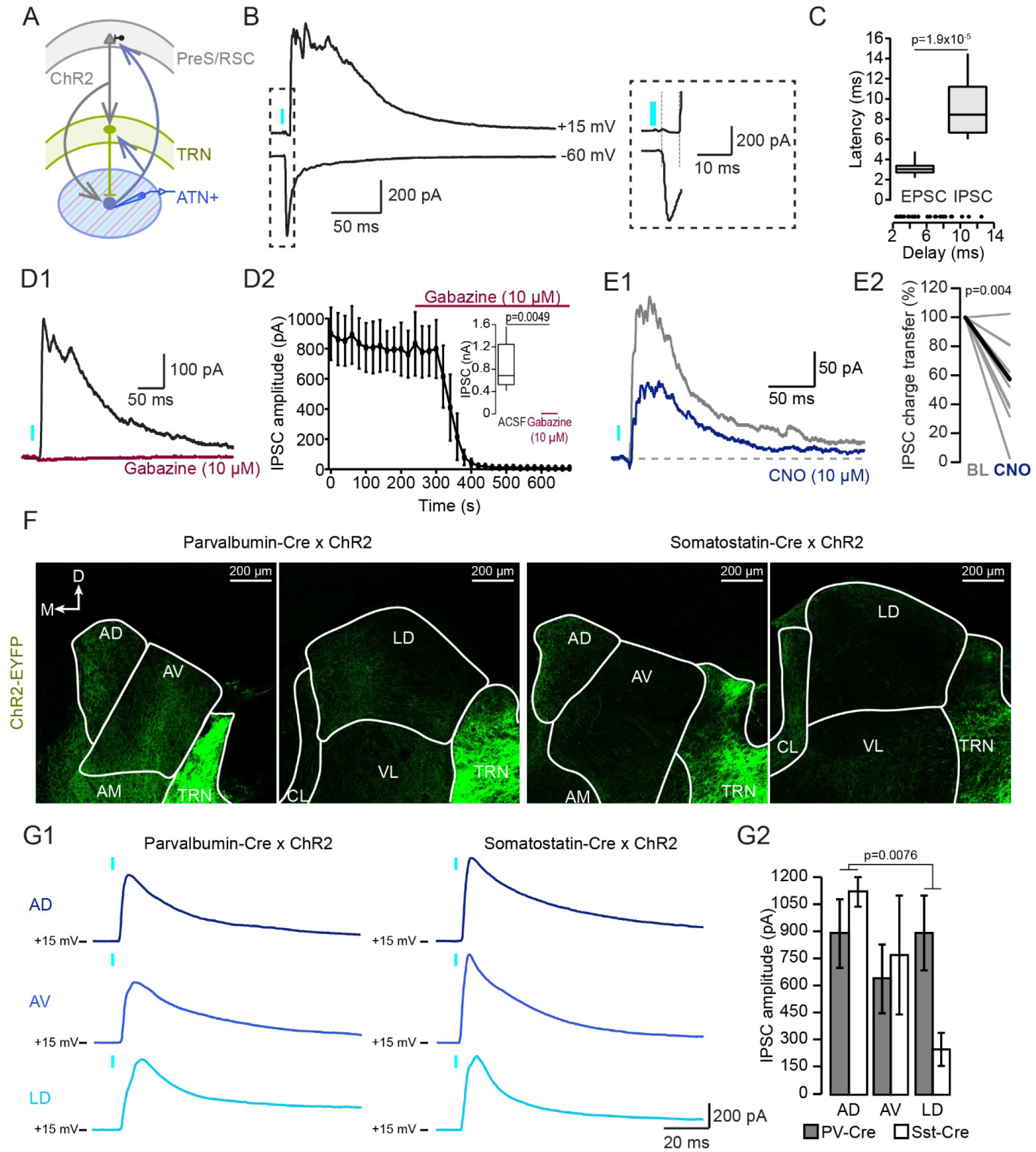
PreS/RSC afferents mediate feedforward inhibition onto anterior thalamus through recruiting burst discharge in PV- and Sst-expressing TRN cells. **(A)** Simplified scheme of the hypothesized circuit studied *in vitro*. **(B)** Typical current responses of an ATN+ neuron held successively at -60 and +15 mV to record EPSC and IPSC. Portion indicated by dotted rectangle is expanded on the right. Response latencies (grey vertical lines) were measured from LED onset. **(C)** Top: Box-and-Whisker plot of EPSC and IPSC latencies in ATN+ neurons (n = 24). Bottom: Delays between the onset of the IPSC and the EPSC for all experiments. Wilcoxon signed rank-test. **(D1)** A typical ATN+ IPSC before (black) and after (red) bath-application of the GABAA receptor antagonist gabazine. **(D2)** Time course of gabazine action (n = 6). Inset: Box-and-Whisker plot of steady-state IPSC amplitude in ACSF and gabazine (paired Student’s *t* test). **(E1)** IPSC evoked in an ATN+ neuron of a VGAT-Ires-Cre mouse expressing hM4D in anterodorsal TRN. IPSCs measured before (grey) and after (blue) 10 μM CNO bath-application. **(E2)** Representation of the charge transfer of IPSCs in ATN+ neurons (n = 10). Wilcoxon signed rank-test. **(F)** Confocal micrographs of ChR2-expressing PV-Cre (left) and Sst-Cre (right) coronal brain sections of ATN+. **(G1)** Representative IPSCs elicited in ATN+ neurons held at +15 mV. **(G2)** Histogram of IPSC amplitudes in AD (n = 6 for both PV- and Sst-Cre mice), AV (n = 6 for both) and LD (n = 6 for both). 2-factors ANOVA with factors ‘nucleus’ and ‘cell type’, p = 0.036 for ‘nucleus’, p > 0.05 for ‘cell type’, post hoc Student’s *t* test with Bonferroni correction: α = 0.017 for IPSC amplitude between nuclei regardless of cell type.

The TRN contains subnetworks of PV- or Sst-expressing cells with possibly different functions (Clemente-Perez et al., 2017). We determined the contribution of these subnetworks to ATN+ inhibition using PV-Cre and Sst-Cre mice expressing ChR2 in anterodorsal TRN. ChR2-positive fibers were visible throughout the AD, AV and LD in both mouse lines (Fig. 4F), and rapid IPSCs were elicited by activation of both PV- and Sst-expressing TRN cells in all thalamic nuclei (Fig. 4G1,G2), suggesting a contribution of both subnetworks to ATN+ inhibition.

### Anterodorsal TRN is implied in the tuning of HD cells and contributes to PreS/RSC-induced action potential discharge dynamics in ATN+

To study the consequences of PreS/RSC inputs on unit activity of ATN+, we performed *in vivo* single unit recordings in freely behaving mice while optogenetically activating PreS/RSC (Fig. 5A-E). We recorded a total of 77 units in ATN+ (n = 7 mice), out of which 18 were HD units (Fig. 5C), as determined according to standard criteria (Yoder and Taube, 2009) (Suppl. Fig. 4A-C). Firing patterns of single units (n = 61/77 responsive units in ATN+, amongst which 14/18 HD units), analyzed through raster plots, peri-event histograms and z-score analysis of the changes in firing rate compared to baseline (Fig. 5A,B, Methods), fell into 4 distinct classes (Fig. 5D, Suppl. Fig. 4). A first major class (n = 26) showed a mixed response composed of an early increase in firing rate that was followed by a decrease and, occasionally, a late increase in firing, reminiscent of a rebound response (Fig. 5A,B). The three other classes showed one of these three response components in isolation (Suppl. Fig. 4). A second small class (n = 3) showed the initial rapid increase in firing rate only. A third large class (n = 18) primarily showed a decrease in firing rate, whereas a fourth class (n = 7) showed only a late increase in firing rate. Time-wise, early increases in firing rate showed an onset latency (relative to the start of the light pulse applied through the optic fibers implanted over PreS/RSC) that was markedly shorter than the decreases and the late increases in firing rate (Fig. 5E). Importantly, HD units responded with similar discharge patterns to PreS/RSC stimulation (Fig. 5D), showing notably decreases in firing rate (8/14 units) and mixed responses (3/14 units). This classification shows that PreS/RSC inputs efficiently control ATN+ discharge dynamics with a consistent temporal pattern. Importantly, a substantial fraction of these units responded with transient firing decreases, including also HD units. This pattern is consistent with an inhibitory synaptic input, implying a possible recruitment of anterodorsal TRN, or of other, still unidentified inhibitory feedforward projections.

**Figure 5.**
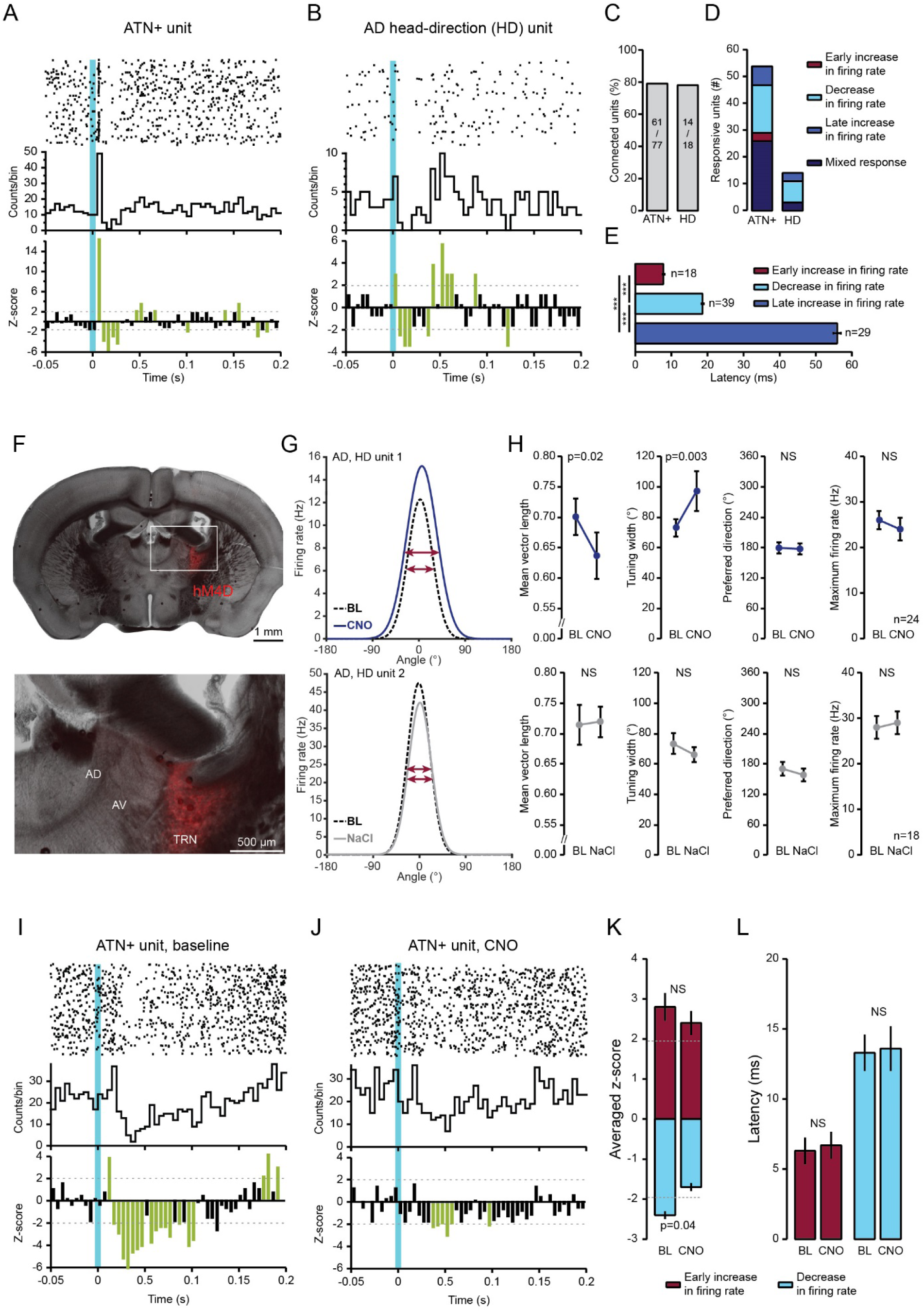
Anterodorsal TRN is implied in the tuning of HD cells and contributes to PreS/RSC-induced action potential discharge dynamics in ATN+. **(A, B)** Raster plot, cumulative histogram and z-score analysis for two characteristic unit responses. Blue vertical line denotes time of light application. Z-score bins labeled in green highlight significant changes in firing rate. **(C)** Total number of light-responsive ATN+ and HD-tuned units. **(D)** Stacked bar graph showing number of units grouped into four classes based on their response properties. See Suppl. Fig. 4 for further examples. **(E)** Histogram of response latencies. Mann-Whitney U tests and Bonferroni correction: α = 0.017 **(F)** Sections for the anatomical verification of silicone probe implantation in VGAT-Ires-Cre mice expressing the chemogenetic silencer hM4D-mCherry (red) in anterodorsal TRN. **(G)** Examples of HD tuning curves for two units prior to and after injection of CNO or NaCl. Horizontal arrow shows tuning width. **(H)** Quantification of the changes in tuning parameters by CNO or NaCl. Far left: Rayleigh vector length size. Middle left: Width of tuning curve. Middle right: Preferred direction. Right: Firing rate at preferred direction. See Methods for details on analysis and Suppl. Fig. 5 for more examples. Paired Student’s *t* tests for Rayleigh vector length size and preferred direction, Wilcoxon signed rank-test for width of tuning curve and firing rate at preferred direction **(I, J)** Same as A, for a unit before (I) and after CNO (J). (**K**) Mean z-scores for early increases (red bar) and decreases (blue bar) in firing rates prior to and after CNO injection. Paired Student’s *t* tests. (**L**) Latencies from photostimulation for the same units as in (K), Mann-Whitney U tests.

Chemogenetic silencing of anterodorsal TRN was used to directly evaluate the role of TRN inhibition for HD units specifically and for ATN+ firing dynamics more generally.

First, we recorded HD units with silicone probes targeted stereotaxically to the AD, the site of HD cells (5 mice, Fig. 5F) (Taube, 1995). We compared the tuning, tuning width, preferred direction and firing rate at the preferred direction of the HD units during a baseline session and 40 min after i.p. injection of CNO (1 – 2 mg/kg) (n = 24) or NaCl (n = 18) (Fig. 5G,H, Suppl. Fig. 5), thereby ensuring that unit properties remained unaltered (see Methods). Overall, we found a decrease in tuning and an increase in the tuning width after CNO injection compared to baseline, whereas preferred direction and maximal firing rate remained unaltered. NaCl injection did not induce changes in any of these four parameters. Therefore, the baseline tuning curve of HD cells in AD in freely moving conditions deteriorates upon a decrease in TRN activity.

We also chemogenetically silenced anterodorsal TRN while recording from a subset of ATN+ units responding to PreS/RSC photostimulation with a decrease in firing rate (n = 15, 8 HD units, 7 untuned units) or with an early increase in firing rate (n = 10, 3 HD units, 7 untuned). These units were recorded in three VGAT-Ires-Cre mice expressing hM4D in the TRN and ChR2 in the PreS/RSC. Following a baseline recording session, mice were recorded again 40 min after CNO i.p. injection (Fig. 5I,J, see Methods). After chemogenetic silencing of the TRN, the mean z-score, calculated from onset to offset of significant decrease in firing rate, was significantly reduced (Fig. 5K). The latency of decrease in firing rate was not altered in the 11 units in which PreS/RSC stimulation still evoked a significant decrease in firing rate in CNO (Fig. 5L). I.P. injection of CNO did not alter the averaged z-score of the light-evoked early increase in firing rate nor the latency of these events (Fig. 5K,L). These data suggest that the feedforward decrease in firing rate observed in ATN+ units upon PreS/RSC activation is at least partially mediated by the anterodorsal TRN.

### Anterodorsal TRN inhibition biases navigational search strategies in the Morris water maze

To probe the role of anterodorsal TRN in spatial navigation, we chose the hidden platform version of the Morris water maze (MWM). In this maze, both ATN-dependent and visual cue-dependent allocentric navigational strategies were reported (Garthe and Kempermann, 2013; Stackman et al., 2012). Mice were trained over 10 days to learn the hidden platform in a maze surrounded by visual landmarks, followed by a 10-day reversal learning during which the platform was located in the opposite quadrant (Suppl. Fig. 6A). We hypothesized that chemogenetic suppression of anterodorsal TRN activity, and reduction of PreS/RSC-mediated ATN+ inhibition, would alter navigational behavior once the animal had to rely on HD-dependent strategies. We also asked whether there was a bias in search strategies already in the course of spatial learning.

**Figure 6.**
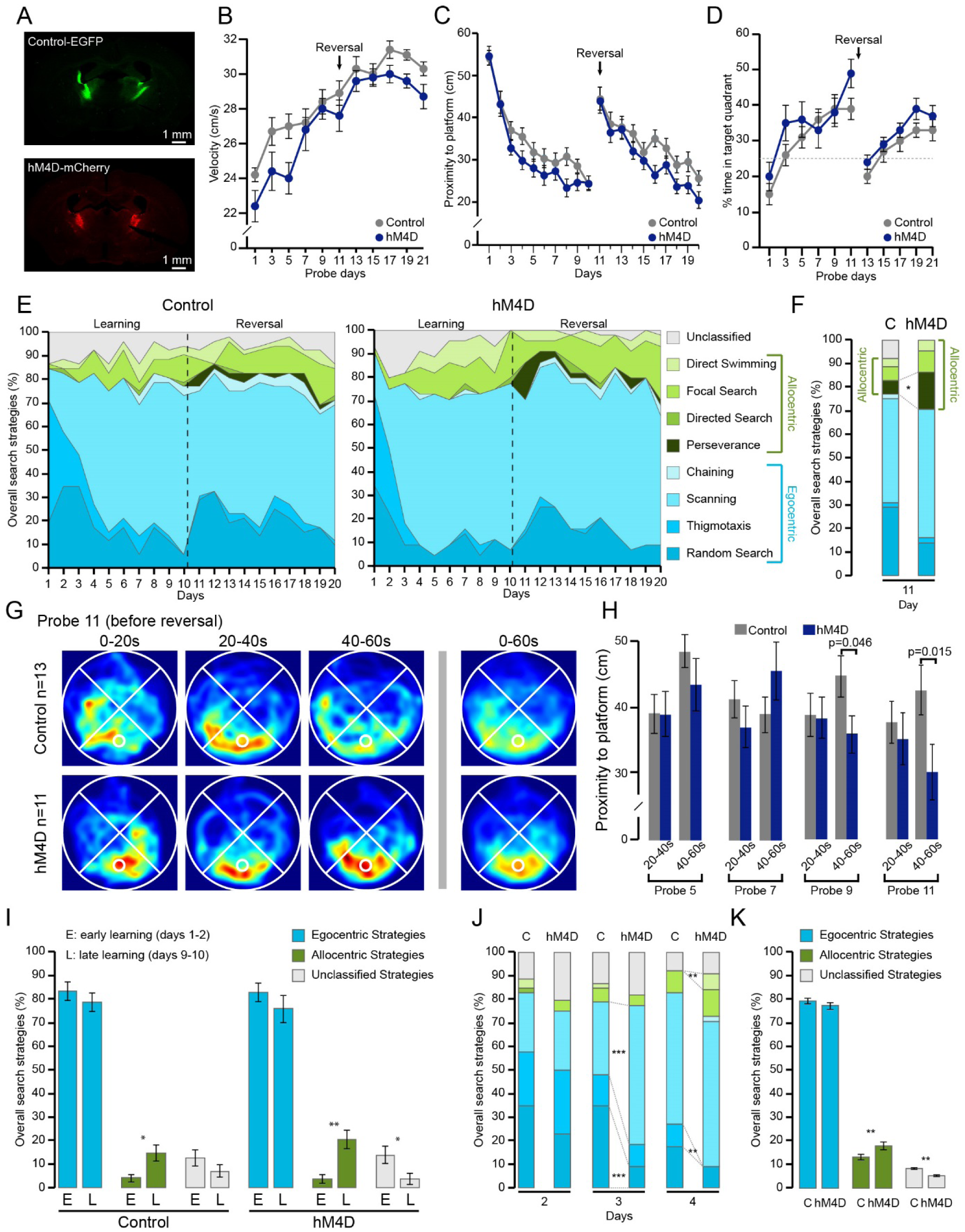
Anterodorsal TRN inhibition biases navigational search strategies in the Morris water maze. **(A)** Epifluorescent micrographs of VGAT-Ires-Cre mouse coronal brain sections at Bg -0.8 mm for a control mouse (top) and a test mouse (bottom). Color codes indicate expression products. **(B)** Mean swim velocities of control (n = 13) and hM4D (n = 11) mice during probe sessions. 2-factors RM ANOVA with factor ‘day’ and ‘condition’, p < 2×10^−16^ for ‘day’ and p = 0.04 for ‘condition’. Post hoc Student’s *t* tests with Bonferroni correction: α = 0.005 for ‘condition’: not significant. **(C)** Graph of the mean proximity to the platform during training sessions. 2-factors RM ANOVA with factor ‘day’ and ‘condition’, p < 2×10^−16^ for ‘day’ and p = 0.04 for ‘condition’. Post hoc Student’s *t* tests and Mann-Whitney U tests with Bonferroni correction: α = 0.003 for ‘condition’: not significant. **(D)** Graph of the percentage of time spent in the target quadrant during probe sessions. Chi-square test against 25% chance, significant at days 1, 7, 9, 11 for Control and days 5, 9, 11, 19, 21 for hM4D. **(E)** Stacked area graphs of search strategies used by control (left) and hM4D (right) mice during trial sessions. **(F)** Proportion of overall strategies at day 11 for control (C) and hM4D (D) mice. Chi-square tests for ‘allocentric strategy’ and for ‘Perseverance’, p < 0.05 for both. **(G)** Time-binned (20 s bins) and overall average occupancy plots during the last probe session of the learning phase. Hot colors indicate greater occupancy and are equally calibrated in all plots. **(H)** Histogram of the mean proximity to the platform of control and hM4D mice during binned-probe sessions. Student’s *t* test for Control vs hM4D at late time bin (40 – 60 s). **(I)** Averaged proportion of egocentric, allocentric and unclassified strategies used during the early (E, days 1 and 2) and late (L, days 9 and 10) learning phase. Wilcoxon signed rank-tests, p = 0.02 for allocentric strategies in control mice, p = 0.008 and p = 0.03 for allocentric and unclassified strategies in hM4D mice, respectively. **(J)** Proportion of overall strategies at day 2, 3 and 4 for control (C) and hM4D (D) mice. Chi-squared tests for ‘Scanning’ and for ‘Random Search’ at day 3, p < 0.001 for both. Chi-square tests for ‘Direct Swimming’ and for ‘Thigmotaxis’ at day 4, p < 0.01 for both. **(K)** Averaged proportion of egocentric, allocentric and unclassified strategies used during the whole experiment. Student’s *t* test comparing Control (C) vs hM4D (D).

We used two groups of mice: “control” VGAT-Ires-Cre and “hM4D” VGAT-Ires-Cre mice that expressed non-hM4D related proteins or the hM4D specifically in the anterodorsal TRN, respectively (Fig. 6A, Suppl. Fig. 7). In each of the four daily test sessions, entry points into the maze were randomized across the quadrants to enforce the use of allocentric strategies. Both groups became faster swimmers in the course of the task and showed no significant difference in their mean swimming velocity during the 60-s probe sessions (with platform removed), although there was a light trend for control mice to be faster (Fig. 6B). We thus analyzed the proximity to the platform instead of the latency to platform to account for possible effects of differences in swim speed (Awasthi et al., 2019). Based on this measure, both groups performed similarly, as indicated by a comparable decrease of the mean proximity to the platform during test sessions (Fig. 6C). Moreover, from days 5 – 7 of training, the percentage of time spent in the target quadrant was above chance level for both groups during probe sessions (Fig. 6D) These results show an overall comparable, if not slightly better performance of hM4D mice, but they do not provide information about the navigational strategies used. We hence classified swim trajectories on a trial-by-trial basis for all test sessions according to previously described criteria for allo- and egocentric strategies (Suppl. Fig. 6B-D) (Garthe and Kempermann, 2013; Rogers et al., 2017). Figure 6E shows that mice use a mix of trajectories reflecting the use of both ego- and allocentric strategies (Fig. 6E,I,K). Focusing first on early phases of reversal learning (day 11), hM4D mice showed perseverance around the previous platform location, while control animals reverted to trajectories consistent with egocentric strategies (Fig. 6F). If perseverance was indeed reflecting a decreased ability to change navigational strategy once the correct platform location was learned, signs of perseverance should also be seen in the course of learning. Indeed, when inspecting time-binned occupancy plots during probe sessions, hM4D mice persevered searching at the platform position for the whole 60 s-probe session, whereas control mice shifted to a dispersed search pattern of other regions of the pool during the last 20 s of the session. This was particularly the case during the last 2 probe sessions of the learning (beginning of day 9 and 11) (Fig. 6 G,H).

Inspired by the finding on the hM4D mice’s possibly compromised ability to deploy egocentric strategies during reversal learning, we asked whether evidence for biased strategy selection could also be found during initial platform learning. As is characteristic for the MWM, there was an increase in the proportion of allocentric strategies from day 1-2 to day 9-10 in both control and hM4D mice (Fig. 6 I,J) (Garthe and Kempermann, 2013). However, hM4D mice did so in temporal anticipation, showing significantly more scanning and less random search on day 3, and more direct swimming and less thigmotaxis on day 4 (Fig. 6J). hM4D mice also used an overall greater proportion of allocentric strategies across both learning and reversal learning than control mice (Fig. 6K). Together, suppression of anterodorsal TRN activity 1) alters navigational behavior at reversal learning and 2) biases the search patterns towards allocentric strategies during initial learning.

## Discussion

Anatomical and physiological identification of synaptic inputs to TRN has repeatedly opened a novel point of view for the TRN’s active role in gating sensory information flow to and from the cortex (for review, see (Crabtree, 2018)). We uncover here a previously undescribed excitatory input to TRN from the parahippocampal dPreS and the multisensory-associative RSC. This dPreS/RSC input recruits feedforward TRN inhibition to ATN and to HD-tuned neurons in AD. We thus identify a novel corticothalamic loop that expands the TRN’s gating function to the domain of head-orientation signals. Our behavioral experiment indicates that this pathway offers a possible synaptic mechanism contributing to the flexible selection of navigational strategies during spatial navigation. More generally, we favor a view of the TRN as a multi-modal saliency selector that interfaces between acute cognitive demands, such as attentional switching or spatial re-orientation, and the recruitment of the appropriate sensory and HD signals.

Retrograde tracing from the anterodorsal portion of the TRN labeled several prefrontal cortical afferents that were described previously (Cornwall et al., 1990; Dong et al., 2019; Lozsádi, 1994). Our novel observation is a continuous band of TRN afferents along the presubicular-retrosplenial axis that starts at the border from subiculum to the PreS, thus at the transition from the three-layered hippocampal to the six-layered parahippocampal complex. These projections arise in deep layers of PreS and RSC, to where projections to ATN were previously retrogradely traced (Wright et al., 2010). Anterodorsal TRN may additionally receive inputs from the centrolateral thalamic nucleus (Krout et al., 2001) and from lateral hypothalamus (Herrera et al., 2016). The functional diversity of these afferents points to the anterodorsal TRN as an assembly of highly integrative neurons that gate thalamic activity appropriately for particular sensory and attentional navigational behaviors.

We note here that both RSC and dPreS target LD and AD preferentially, while projections to AV are minor. AD and LD are thought to functionally cooperate within the HD system (Perry and Mitchell, 2019; Simonnet and Fricker, 2018), possibly acting as first- and higher-order nucleus, respectively (Peyrache et al., 2019). The AV, together with the anteromedial nucleus, has been so far associated with a theta-generating system innervated by vPreS (Perry and Mitchell, 2019). The limited spatial resolution of our tracing methods does not currently allow to verify whether anterodorsal TRN is also subdivided into sectors corresponding to this functional subdivision of ATN+. Moreover, the detailed contributions of presubicular, and of the granular and agranular portions of RSC, warrant further investigation. More work is also required to elucidate the detailed organization of the synaptic connectivity from PreS/RSC to anterodorsal TRN and from there to ATN. Interestingly, single-cell labeling identified rat anterodorsal TRN cells with axons bifurcating to innervate both AD and LD (Pinault and Deschênes, 1998). AM-projecting TRN cells were located more ventrally. Anterodorsal TRN may thus contain cells jointly innervating AD and LD, further substantiating a shared function.

Our characterization of a cortical excitatory innervation of TRN by PreS/RSC-excitatory input reveals a combination of commonalities but also notable differences to the canonical form of cortical input to sensory TRN that arises from layer 6 corticothalamic neurons of corresponding primary cortex (Usrey and Sherman, 2019). Layer 6 synapses on TRN cells show a high glutamate receptor content (Golshani et al., 2001), high unitary amplitude (Golshani et al., 2001), faster rise and decay times (Gentet and Ulrich, 2004), smaller NMDA/AMPA ratios (Astori and Lüthi, 2013) and marked PPF (Astori and Lüthi, 2013; Castro-Alamancos and Calcagnotto, 1999; Crandall et al., 2015) compared to their thalamocortical counterparts. The presence of PPF is an important criterion to classify layer 6 corticothalamic afferents as modulators rather than drivers (Sherman and Guillery, 1998). While dPreS/RSC-TRN synapses are comparable in terms of unitary amplitude, NMDA/AMPA ratio and EPSC waveform, there is a prominent lack of PPF at dPreS/RSC afferents and a moderate entrainment of firing during repeated stimulation. Rather than being modulators, the PreS/RSC inputs thus share a short-term plasticity profile reminiscent of the driver inputs that count as the principal information-bearing synapses. Top-down driver input is so far known for corticothalamic layer 5 projections to higher-order thalamic nuclei that show a number of morphological hallmarks (Usrey and Sherman, 2019). A driver profile, as we suggest here for the first time for a TRN afferent, implies that anterodorsal TRN conveys direct system-relevant information that is faithfully transmitted to its projection targets. We cannot exclude, however, that PreS and RSC afferents, if stimulated separately, would show different short-term plasticity including PPF and arise from different cortical layers (Simmonet and Fricker 2018). A further noteworthy point is that both PV+ and Sst+ TRN neurons innervate the AD, AV and LD with comparable strength, pointing to functional differences compared to sensory first-order thalamic nuclei (Clemente-Perez et al., 2017).

When PreS/RSC afferents were activated *in vivo*, a majority of ATN+ units (∼79%) and of HD units (77%) responded with changes in action potential firing. We observed sequential firing rate increases and decreases that fall within the time ranges expected for a feedforward inhibition (Crandall et al., 2015). These firing dynamics could result from a mix of afferent mono- and polysynaptic activities because excitatory PreS/RSC neurons project widely to cortical, hippocampal and subcortical targets (Clark et al., 2018; Mitchell et al., 2018; Simonnet and Fricker, 2018), including to the lateral mammillary bodies (Yoder et al., 2015). Potential long-range inhibitory projections that could have been virally transformed also cannot be excluded. However, in strong support of a recruitment of TRN, we show that decreases in firing rate in ATN+ and HD units are attenuated once anterodorsal TRN is silenced while all increases in firing rate are preserved. Moreover, the latencies of the initial firing increases and decreases were short and showed little variability from unit to unit, which fits better with a monosynaptic feedforward inhibitory circuit, such as the one recruited by direct TRN activation through PreS/RSC, rather than a polysynaptic one.

What could be the functional consequences of PreS/RSC-mediated feedforward inhibition? The precision tuning of the thalamic HD cells is essential for the cortico-hippocampal representation of space (for review, see (Peyrache et al., 2019)). It is distinctly sharper than in the upstream lateral mammillary body (Blair et al., 1998), suggesting that active inhibition is at work at thalamic levels to suppress firing at angles outside the preferred direction. TRN is a likely candidate for this inhibition, because, as we show, the HD tuning of AD cells degrades with a silenced TRN. As the effect size we measure is comparable to the one observed after PreS lesion (Goodridge and Taube, 1997), PreS-dependent TRN regulation could act in a circuit analogous to that of descending corticothalamic control of thalamic sensory receptive fields (Temereanca and Simons, 2004). Sharpening could also arise via recurrent synaptic activity between TRN and AD to coordinate activity in similarly tuned HD neurons (Peyrache et al., 2015). Once important visual landmark information has to processed by the HD system, PreS/RSC could act through TRN to enforce AD discharge in response to novel cues, thereby helping to reset the tuning via generating inhibition-rebound firing. Consistent with this disynaptic circuit, it has been shown that updating of HD tuning occurs rapidly (< 100 ms), which requires short synaptic delays (Zugaro et al., 2003). Moreover, there is now first evidence that acute optogenetic inhibition of TRN indeed prevents the updating of HD units to rapidly changing environments (Duszkiewicz et al., 2019). TRN-driven ATN bursting might also be an important component in oscillatory patterns observed within ATN+, such as the one proposed to occur in AD (Peyrache et al., 2019; Peyrache et al., 2015) or in AV (Tsanov et al., 2011), which are probably relevant for linking spatial information to hippocampal memory processing.

To date, behavioral evidence for a role of the HD system in behavioral navigation is limited (Butler et al., 2017; Taube et al., 1992; Valerio and Taube, 2012; van der Meer et al., 2010) and the role of specific HD circuits, including dPreS and RSC, just starts to be behaviorally explored. Experimental effort typically targets egocentric strategies, for example by moving the MWM relative to landmarks between trials (Stackman et al., 2012), by studying navigation in darkness (Yoder et al., 2019) or by putting rats upside down (Calton and Taube, 2005). In contrast, the idea that spatial navigation requires an on-going switching between a range of possible strategies has not been much pursued, although it is known for human studies (Miniaci and De Leonibus, 2018). For example, a recent study using a hippocampus-specific synaptic knockout animal interpreted perseverant behavior at the platform location as a lack of forgetting (Awasthi et al., 2019), but the question of possible bias in navigational strategies was not addressed. Trajectory analysis in the MWM thus offers itself as an interesting approach to follow on the evolution of navigational behavior under well-controlled landmark conditions while simultaneously allowing egocentric strategies (Dolleman-van der Weel et al., 2009; Garthe and Kempermann, 2013). We found a preferential use of allocentric strategies when anterodorsal TRN was suppressed, suggesting that egocentric navigation was less efficient. This is reminiscent of an ATN lesion study (Stackman et al., 2012) and goes along with preserved visual and motor behaviors of the swimming animals. Therefore, it is also unlikely that TRN-dependent inhibitory effects on nuclei other than ATN, such as on intralaminar nuclei to attenuate escape responses, (Dong et al., 2019), are primary causes for the change in search strategies. Anterodorsal TRN activity seems to be critically required at moments when there is a mismatch between allocentric cues and new platform location, such that novel relations between external landmarks and navigational strategies, which depend on HD cells, have to be built. Interestingly, the RSC has been proposed as an area involved in allocentric navigation and memory formation, but also in the switching between allo- and egocentric strategies to optimize navigational goals (Mitchell et al., 2018). In particular, its strong connections to limbic thalamus have been implied in the solving of spatial problems (Clark et al., 2018). Similar more complex roles in spatial navigation have recently been proposed for dPreS (Yoder et al., 2019), which has been primarily analyzed as part of the hierarchy of the egocentric coding system (Dumont and Taube, 2015; Peyrache et al., 2017; Taube et al., 1990). Our work does not currently disentangle between the distinct roles of these two brain areas. However, it has managed to pinpoint to the existence of a possibly fine switching mechanism at the interface between major brain areas that, when perturbed, preserved overt navigational performance but compromised it at challenging moments that could pose existential threats.

This work integrates TRN function into the brain’s control of sensory-guided spatial navigation. The anterodorsal sector of TRN, located at the *limbus*, the ‘edge’ of the TRN, is a site of complex integration where navigational, attentional, motor and emotional information may be combined for precise control of anterior thalamus-dependent navigational systems. As a perspective arising from this work, we suggest that neuropsychological screening for deficits in navigational flexibility may be useful in the diagnosis of disorders linked to TRN dysfunction, such as in neurodevelopmental disorders linked to attentional deficits (Krol et al., 2018) and in schizophrenia (Wilkins et al., 2017).

## Acknowledgements

We thank Andreas Lüthi for providing training for the multiwire recordings, Adrien Peyrache for providing training for the silicone probes, Cyril Herry for helping with single unit sorting and analysis, Leonardo Restivo for helping with the design of the Morris water maze experiment and careful reading of the manuscript, Christian Lüscher for providing the VGAT-Ires-Cre mouse line, Desdemona Fricker, John Huguenard, Ralf Schneggenburger for constructive discussions, Simone Astori for critical reading of the manuscript, all lab members for constructive input for the manuscript and discussions in the course of the project.

## Author contributions

GV carried out and analyzed all experimental tracing, *in vitro, in vivo* and behavioral data, and also contributed to the design of the *in vivo* and behavioral experiments. ZR gained first evidence for the anatomical connectivity between dPreS/RSC and anterodorsal TRN and initiated the *in vivo* unit recordings. RC wrote the Matlab code for the analysis of head-direction data and MWM Strategy. EB contributed to viral injections and the MWM experiments, GK to the antero- and retrograde tracing data. AG carried out the *in vitro* recordings in PV- and Sst-Cre mice. VP assisted with anatomical analysis. LMJF contributed to the surgery and analysis for *in vivo* experiments. AL supervised the project and wrote the manuscript with contribution of GV.

## STAR Methods

### Animal husbandry and ethical approval

We used mice of either sex from the C57BL6/J line and from the Slc32a1^tm2(cre)Lowl^ line, commonly referred to as VGAT-Ires-Cre line (Jackson Labs, generated by Dr. B. Lowell, Beth Israel Deaconess Medical Center, Harvard) (Vong et al., 2011), male C57Bl/6J;129P2_Pvalbtm1(cre)Arbr/J mice, referred to here as PV-Cre mice, and male B6N.Cg-Sst<tm2.1(cre)Zjh>/J mice, referred to here as Sst-Cre mice. These three transgenic lines express the Cre-recombinase either in VGAT-, PV- or Sst-positive neurons, respectively. All animals were housed in a temperature and humidity-controlled animal house with a 12/12 h light-dark cycle (lights on at 9 a.m.) and water and food available *ad libitum.* The VGAT-Ires-Cre line was originally generated on a mixed C57BL/6;FVB;129S6 genetic background and backcrossed to C57BL6 ever since. PV-Cre and Sst-Cre lines were maintained on a C57BL6 background. VGAT-Ires-Cre and PV-Cre were used as homozygous, whereas the Sst-Cre mice were heterozygous. For anatomical tracing (retrograde and anterograde), mice (n = 24) were transferred into a housing room with similar conditions on the day prior to injection. They remained there for 7 days after injection before perfusion and tissue processing. For viral injections, mice were transferred into a P2 safety level housing with similar conditions on the day prior to the injection. They remained there 3 – 5 weeks before being used for *in vitro* electrophysiology (n = 57), 2 – 3 weeks before surgical implantation for *in vivo* electrophysiology (n = 9), and 2 – 3 weeks before behavioral experiments (n = 24, only males). All experimental procedures complied with the Swiss National Institutional Guidelines on Animal Experimentation and were approved by the Swiss Cantonal Veterinary Office Committee for Animal Experimentation.

### Anatomical tracing and verification of recording and injection sites

#### Retrograde tracing

C57BL6/J mice, 4-8-week-old, were anesthetized with 5 % isoflurane and fixed onto the stereotaxic frame. During the surgery, the anesthesia level was reduced to 1-3 % and N_2_O was added if the surgery lasted > 1 h. Analgesia was ensured through Carprofen (5 mg/kg i.p.). Craniotomies were performed above the sites of injection at (anteroposterior (AP), mediolateral (ML), depth from cortical surface (DV), in stereotaxic coordinates from Bregma): -0.7, ±1.5, -3.1 to target the anterodorsal TRN. Glass pipettes (5-000-1001-X, Drummond Scientific, Broomall, PA) were pulled on a vertical puller (Narishige PP-830, Tokyo, Japan) and backfilled by capillarity with fluorescent latex microspheres (Red Retrobead™, Lumafluor). Using a Picospritzer III, pressurized air pulses (15 psi, 10 ms) were applied every 10 s for 10 min to inject the retrobeads. After 4 – 7 days, mice were perfused and their brains collected for immunostainings.

#### Anterograde tracing

The anesthetic and surgical procedures were the same as the ones used for retrograde tracing. The coordinates of injection were (AP, ML, DV): -3.8, ±1.6, -1.0 for RSC, -3.8, ±2.3, -1.6 for PreS. Glass pipettes were backfilled by capillarity with the plant lectin anterograde tracer *Phaseolus vulgaris-*leucoagglutinin (PHAL-L, Vector Laboratories, Cat. No. L-1110). PHAL-L was chosen as it permits focal labeling with little spread, which seemed appropriate to target PreS and RSC as specifically as possible. A chlorinated silver wire was inserted into the pipette and a reference electrode attached to the mouse tail. The PHAL-L was electroporated with a 5-μA current, 7 s on/off loop for 20 min, applied with a home-made current isolator and a Master-8 (Master-8 Pulse Stimulator, A.M.P.I., Jerusalem, Israel). After 5 – 7 days, mice were perfused and their brains collected for immunostainings.

#### Perfusion and tissue processing

Mice were injected i.p. with a lethal dose of pentobarbital. Intracardial injection of ∼45 ml of paraformaldehyde (PFA) 4 % was done at a rate of ∼2.5 ml/min. Brains were post-fixed in PFA 4 % for at least 24 h at 4°C. Brains were sliced with a Vibratome® (Microtome Leica VT1000 S, section thickness: 100 μm, speed: 0.25-0.5 mm/s and knife sectioning frequency: 65 Hz) in 0.1 M phosphate buffer (PB). Brain sections were either directly mounted on slides or disposed in twelve-well plates filled with 0.1 M PB for immunohistochemistry.

#### Immunofluorescent labeling

100 μm-thick brain sections were washed 3 times in 0.1 M PB and transferred to a blocking solution containing 0.1 M PB, 0.3 % Triton, 2 % normal goat serum (NGS) for 30 min. The first antibody solutions also contained 0.1 M PB, 0.3 % Triton, 2 % NGS. For PHAL-L injected mice, we added 1:8000x of rabbit anti-PHAL-L (Vector Laboratories, AS-2300, RRID: AB_2313686) and 1:4000x of mouse anti-PV (Swant, PV235, RRID: AB_10000343). For retrobead-injected and virally injected PV-Cre and Sst-Cre mice, we added 1:4000x of mouse anti-PV (Swant, PV235, RRID: AB_10000343). Sections were kept at 4°C for 48 h on a shaking platform. After 3 washings in 0.1 M PB, we added a secondary antibody solution containing 0.1 M PB, 0.3 % Triton, 2 % NGS and, when appropriate, 1:500x of goat anti-rabbit Cyanine Cy3^TM^ (Jackson Immunoresearch, 111-165-003, RRID: AB_2338000), 1:500x of goat anti-mouse Cy5^TM^ Jackson Immunoresearch, 115-175-146, RRID: AB_2338713) and/or 1:500x of goat anti-mouse Alexa Fluor® 488 (Jackson Immunoresearch, 115-545-003, RRID: AB_2338840). Sections were mounted on slides and covered with a mounting medium (Vectashield).

300 μm-thick brain sections obtained from patch-clamp recording sessions were post-fixed in 4 % PFA for at least 24 h. Brain sections were washed 3x in 0.1 M PB and then pretreated with a solution containing 0.1 M PB and 1 % Triton for 30 min. The blocking solution was 0.1 M PB, 1 % Triton, 2 % NGS and was applied for 30 min. The first antibody solution contained 0.1 M PB, 1 % Triton, 2 % NGS, 1:4000x mouse anti-PV (Swant, PV235, RRID: AB_10000343) and was applied for 5 days at 4°C. The secondary antibody solution contained 0.1 M PB, 0.3 % Triton, 2 % NGS, 1:500x goat anti-mouse CY5, (Jackson ImmunoResearch, 115-175-146, RRID: AB_2338713), 1:8000x Streptavidin ALEXA594 (Jackson ImmunoResearch, 016-580-084, RRID: AB_2337250) and was applied for 24 h at 4°C. Sections were mounted on slides and covered with a mounting medium (Vectashield).

#### Microscopy

Electromicrographs of brain slices were taken with a fluorescent stereomicroscope (Nikon SMZ 25) or a confocal microscope (Zeiss LSM 780 Quasar Confocal Microscope). NIS-Elements 4.5 (Nikon), Adobe Photoshop CS5 and Zen lite 2012 were used to merge images from different channels.

### Viral injections

Mice 3-5-week-old were anesthetized using Ketamine-Xylazine (83 and 3.5 mg/kg, respectively) and placed on a heating blanket to maintain the body temperature at 37°C. An initial dose of analgesic was administrated i.p. at the beginning of the surgery (Carprofen 5 mg/kg). The animal was head-fixed on a stereotactic apparatus equipped with an ear and mouth adaptor for young animals (Stoelting 51925, Wood Dale, IL). The bone was exposed at the desired injection site through a small skin incision. Viruses were injected with a thin glass pipette (5-000-1001-X, Drummond Scientific, Broomall, PA) pulled on a vertical puller (Narishige PP-830, Tokyo, Japan). C57BL6/J mice were injected bilaterally with a virus encoding ChR2 (500 nl of AAV1-CaMKIIa-ChR2(H134R)_eYFP-WPRE-hGH, 10^12^ GC, ∼100–200 nl/min) into the PreS (AP, ML, DV): -3.8, +/-2.5, -1.7. VGAT-Ires-Cre mice were injected uni/bilaterally with 500 nl of AAV1-CamKIIa-ChR2(H134R)_eYFP-WPRE-hGH (1×10^12^ GC, ∼100–200 nl/min) into the PreS and/or uni/bilaterally with a virus encoding hM4D -mCherry (500 nl of AAV8-hSyn-DIO-hM4D(Gi)_mCherry, 6.4×10^12^ GC), or hM4D -IRES-mCitrine (500 nl of ssAAV8/2-hSyn1-dlox-A_hM4D(Gi)_IRES_mCitrine-dlox-WPRE-hGHp(A), 3.1×10^12^ GC) or a control AAV8 encoding a hM4D -unrelated construct (500 nl of AAV8-hSyn-FLEX-Jaws_KGC_GFP_ER2, 3.2×10^12^ GC) in the anterior sector of the TRN (AP, ML, DV: -0.8, ±1.35, -3.1). PV-Cre and Sst-Cre mice were injected into the anterior TRN (AP, ML, DV: -0.8, ±1.35, -3.1) with AAV1-EF1a-DIO-ChR2(H134R)_eYFP-WPRE-hGH (1×10^12^ GC, 500 nl, ∼100–200 nl/min).

### In vitro electrophysiological recordings

#### Slice preparation, solutions and recordings

Brain slice preparation, storage and recordings were performed essentially as described (Fernandez et al., 2018). Adult 8-10-week-old C57BL6/J and VGAT-Ires-Cre mice (3 – 4 weeks post viral injection) were briefly anesthetized with isoflurane and their brains quickly extracted. Acute 300-μm-thick coronal brain slices were prepared in ice-cold oxygenated sucrose solution (which contained in mM: 66 NaCl, 2.5 KCl, 1.25 NaH_2_PO_4_, 26 NaHCO_3_, 105 D(+)-saccharose, 27 D(+)-glucose, 1.7 L(+)-ascorbic acid, 0.5 CaCl_2_ and 7 MgCl_2_), using a sliding vibratome (Histocom, Zug, Switzerland). Slices were kept for 30 min in a recovery solution at 35°C (in mM: 131 NaCl, 2.5 KCl, 1.25 NaH_2_PO_4_, 26 NaHCO_3_, 20 D(+)-glucose, 1.7 L(+)-ascorbic acid, 2 CaCl_2_, 1.2 MgCl_2_, 3 myo-inositol, 2 pyruvate) before being transferred to room temperature for at least 30 min before starting the recording. Slices were placed in the recording chamber of an upright microscope (Olympus BX50WI, Volketswil, Switzerland) and continuously perfused at room temperature with oxygenated ACSF containing (in mM): 131 NaCl, 2.5 KCl, 1.25 NaH_2_PO_4_, 26 NaHCO_3_, 20 D(+)- glucose, 1.7 L(+)-ascorbic acid, 2 CaCl_2_ and 1.2 MgCl_2_. This solution was supplemented in all experiments with 0.1 picrotoxin, 0.01 glycine, with picrotoxin removed for the recordings testing for feedforward inhibition (see Fig. 4). Borders of anterior TRN and ATN+ were visually identified in transillumination using a 10x water-immersion objective. Within a selected nucleus, cells were visualized through differential interference contrast optics a 40x water-immersion objective. Infrared images were acquired with an iXon Camera X2481 (Andor, Belfast, Northern Ireland). Cells were patched using borosilicate glass pipettes (TW150F-4) (World Precision Instruments, Sarasota, FL) pulled with a DMZ horizontal puller (Zeitz Instruments, Martinsried, Germany) to a final resistance of 2.5-5 MΩ. A K^+^-based intracellular solution that contained (in mM) 140 KGluconate, 10 Hepes, 10 KCl, 0.1 EGTA, 10 phosphocreatine, 4 Mg-ATP, 0.4 Na-GTP, pH 7.3, 290–305 mOsm, supplemented with ∼2 mg/ml of neurobiotin (Vector Labs, Servion, Switzerland) was used for comparative measurements of the passive cellular properties (Fig. 2B,2C), for the cell-attached recordings (Fig. 3C) and for all current-clamp recordings (Fig. 3E). A Cs^+^-based intracellular solution containing (in mM) 127 CsGluconate, 10 Hepes, 2 CsBAPTA, 6 MgCl_2_, 10 phosphocreatine, 2 Mg-ATP, 0.4 Na-GTP, 2 QX314-Cl, supplemented with ∼2 mg/ml of neurobiotin, pH 7.3, 290–305 mOsm) was used with all the other voltage-clamp protocols. For these solutions, a liquid junction potential of -10 mV was taken into account for the current-clamp data. Signals were amplified using a Multiclamp 700B amplifier, digitized via a Digidata1322A and sampled at 10 kHz with Clampex10.2 (Molecular Devices, San José, CA).

#### Recording protocols, optogenetic stimulation and analysis

Immediately after gaining whole-cell access, cell resistance (R_m_) and cell capacitance (C_m_) were measured in voltage-clamp at -60 mV through applying 500 ms-long, 10-20 mV hyperpolarizing steps (5 steps/cell). Then the recording was switched to current-clamp to measure the resting membrane potential (RMP). Squared somatic current injections (−50 to -300 pA for 500 ms, 4 injections/cell) hyperpolarized neurons below -100 mV from membrane potentials between -50 to -60 mV and induced repetitive burst discharge in TRN neurons and single burst discharge in thalamic neurons (Fig. 2B, 2C). Squared current injections of increasing amplitude (step size, 50 pA, 500 ms) were used to depolarize the neurons and generate tonic firing. Action potential properties were measured at the rheobase.

Whole-field blue LED (Cairn Res, Faversham, UK) stimulation (455 nm, duration: 0.1 to 1 ms, maximal light intensity 3.5 mW, 0.16 mW/mm^2^) in voltage-clamp (−60 mV) was used to assess the connectivity of TRN and ATN+ neurons through fibers arising from the PreS/RSC. EPSCs were elicited through single light pulses every 20 s, with a 5 mV hyperpolarizing step to control for the access resistance. After a stable baseline of > 2 min, drugs were applied in the bath (40 μM DNQX, 100 μM D,L-APV). To measure NMDA-components, the holding membrane potential was slowly brought to +40 mV where NMDAR-mediated currents were recorded for 2 min before bath-application of D,L-APV. Single light pulses were used in protocols to measure EPSC kinetics and pharmacological properties (Fig. 2D, 2E). The latency from LED onset, EPSC half-width and EPSC weighted decay time constant were measured with Clampfit 10.2. The effect of bath-application of 40 μM DNQX was measured once the reduction of EPSC amplitude reached a steady state. The NMDA/AMPA ratio was measured by dividing the amplitude of the EPSC at +40 mV in DNQX by the amplitude of the EPSC at -60 mV during the baseline and was expressed in percentage.

Minimal stimulation was achieved by progressively reducing the intensity of a single light pulse from its maximum (3.5 mW) to a level where only ∼50 % of the stimuli induced a successful EPSC. The light intensity potentiometer allowing a limited graduation of light intensities, we could include only a few cells (n = 12/50) in which this condition was achieved at 0.28±0.05 mW. In the case of LD neurons, which showed very high amplitude EPSCs with frequent escape currents, none of them reached the criterion to be included. Minimal stimulation was observed for light intensities averaging 0.28±0.05 mW, less than 10 % of the maximum. In a subset of cells (n = 8), we slightly increased light intensity to 0.40±0.08 mW to verify whether failure rate decreased but the amplitude of successful responses was maintained. This was achieved in 5 cells in which failure rate decreased to 0 % but the amplitude of successes was 109±4 % of that found during minimal stimulation, whereas it increased to > 140 % in the remaining 3 cases. All successful EPSCs at minimal light stimulation were visually identified and measured in Clampfit10.2.

Cell-attached recordings of TRN cells (Fig. 3C) were achieved with recording pipettes of ∼5 MΩ resistance, voltage-clamped at 0 mV and ∼0 pA of holding current, while applying single light stimuli at varying light intensity (∼100 stimuli/cell, one every 10 s). Whole-cell access was then established and cells held in current-clamp at their resting membrane potential. Single light stimulations with similar light intensities were given ∼30 times for each cell every 10 s. The number of action currents/action potentials and the interspike interval (ISI) were manually measured on Clampfit10.2. The number of spikes was normalized to the maximum number evoked by the light stimulation. Data were grouped in bins of 0.25 mW of light (Fig. 3C3) and a sigmoidal fit was applied using Igor Pro 7 (WaveMetrics Inc., Lake Oswego, OR). The sigmoidal fit for cell-attached 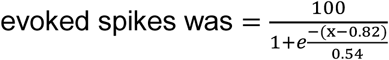. The sigmoid fit for whole-cell 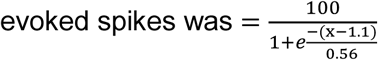.

Paired light stimulations at 1, 2, 5, 10 and 20 Hz were used to assess the short-term plasticity of PreS/RSC-TRN and PreS/RSC-ATN+ synapses. The paired pulse ration (PPR) was expressed as the ratio between the second and the first EPSC amplitude (Fig. 3D). Four responses were elicited for each frequency, with an interval of 20 s between each protocol. The amplitude of EPSCs was measured on the average trace in Clampfit10.2, and traces were not included if spontaneous currents appeared in between the paired stimuli.

For train stimulation, PreS-RSC afferents to TRN and ATN+ neurons were stimulated with 10 light pulses delivered 1/30 s at 2, 5, 8 and 10 Hz while cells were held at -50 to -60 mV in current-clamp (Fig. 3E). Per stimulation frequency and cell, 5 responses were recorded and averaged. Responses were subdivided into sustained (Fig. 3E1) or entrained (Fig. 3E2) responses based on whether or not the first light pulse elicited an action potential. The number of action potentials generated by the train of stimulation was counted on Clampfit10.2. To quantify sustained and entrained responses, the number of action potentials during the 5 first stimulations was compared to the number of action potentials during the 5 last stimulations. In subthreshold responses, the amplitudes of the phasic responses were calculated from the point of positive inflection after a light stimulation to the next positive peak for each of the 10 subthreshold responses. The persistent depolarization was measured as the difference between the baseline value before the train of stimulation and the point of positive inflexion after each light stimulation. The mean persistent depolarization for the last 3 stimulations was used to quantify the steady state response.

To record feedforward inhibitory currents, using the Cs-based intracellular solution defined above, we studied single light-evoked EPSCs recorded in ATN+ cells at -60 mV (uncorrected for a 10 mV junction potential). Then the membrane potential of the cell was slowly brought to +15 mV (uncorrected for a 10 mV junction potential). In 6 ATN+ cells, IPSCs were recorded for 4 min (12 protocols, once every 20 s) for a baseline, then 10 μM gabazine were bath-applied. The amplitude of the IPSCs in gabazine was measured at the steady state. A similar protocol was applied for 10 ATN+ cells recorded in VGAT-Ires-Cre mice expressing the hM4D in TRN cells. Instead of gabazine, 10 μM CNO were bath-applied after the baseline recording of IPSCs. Measures of charge transfer were used to take into account the variable waveform of the IPSCs that were composed of multiple superimposed burst-like synaptic events.

To determine the connectivity of PV- and Sst-expressing TRN cells, brain slices were prepared from PV-Cre and Sst-Cre lines previously injected with ChR2-expressing virus (see above). Using identical recording and light stimulation conditions, evoked IPSCs were quantified in neurons recorded in the different thalamic nuclei AD, AV and LD.

### In vivo single-unit recordings and head-direction monitoring

#### Electrode and fiber preparation

Two types of recording configurations were used. Multi-wire electrodes were implanted for studying response properties of ATN+ to PreS/RSC stimulation. Silicon probes were used for identification and recording of AD HD-tuned units in combination with optogenetic activation of PreS/RSC and/or chemogenetic silencing of the anterodorsal TRN.

Three mice were implanted with the multi-wire electrodes that consisted of 16 individually insulated nichrome wires (13-μm inner diameter, impedance 1 – 3 MΩ; California Fine Wire) contained in a 26-gauge stainless steel guide canula. The wires were attached to a 16-pin connector (CON/16m-V-t, Omnetics) (Courtin et al., 2014), cut at a length of ∼2 mm from the edge of the metal guiding tube and gold-plated using a nanoZ^TM^ device (White Matter LLC, provided by Plexon Inc., Dallas, TX) to a final impedance of 50 – 100 kΩ. A silver wire (Warner Instr.) was soldered to the ground pin of the connector. Six mice were implanted with a single shank linear silicone probe (Neuronexus A1×16-5mm-50-703-Z16 or A1×16-3mm-50-703-Z16).

The optic fibers were built from a standard hard cladding multimode fiber (225 μm outer diameter, Thorlabs, BFL37-2000/FT200EMT), inserted and glued (Heat-curable epoxy, Precision Fiber Products, ET-353ND-16OZ) to a multimode ceramic zirconia ferrule (Precision Fiber Products, MM-FER2007C-2300). The penetrating end was cut at the desired length (∼2 mm) with a Carbide-tip fiber optic scribe (Precision Fiber Products, M1-46124). The other end was polished with fiber-polishing films (Thorlabs). The optic fibers were connected to a PlexBright Optogenetic Stimulation System (Plexon) via home-made patch chord. The connection to the PlexBright Table-top LED Module (Wavelength 465 nm) was achieved through a Mini MM FC 900μm Connector (Precision Fiber Products, MM-CON2004-2300-14-BLK). The other end of the patch chord was inserted into a ceramic zirconia ferrule, fixed with glue and heat-shrinking tube (Allied Electronics, 689-0267) and polished. Before the recording, the patch chord was attached to the implanted optic fiber via a ceramic split sleeve (Precision Fiber Products, SM-CS125S).

#### Surgery

Virally injected C57BL6/J and VGAT-Ires-Cre mice were anesthetized with 5 % isoflurane, fixed on a stereotaxic frame and kept on a feedback-controlled heating pad (Phymep). The level of isoflurane was reduced along the surgery until 1 % and mixed with N_2_O. Craniotomies were opened above the PreS (AP, ML, DV: -3.8, +/-2.5, -1.7 for vertical implantation, -3.8, -3.0, -1.5 with a 30° angle for implantation in diagonal), the left ATN for multi-wire electrodes (AP, ML, DV: -0.8, +0.75, -2.8) or left AD for silicone probes (AP, ML, DV: -0.8, +0.75, -3.3) and the lateral cerebellum with a microdrill (1/005 drill-size). The conjunctive tissue on the skull was removed with a scalpel and the skull was cleaned with iodine-based disinfectant. The skull was then scratched with the tip of the scalpel in a grid-like meshwork of grooves to improve the attachment of the glue (Loctite 401, Koening). Multi-wire electrodes and linear silicone probes were lowered vertically, at approximately 10 μm/s initially and then 1 μm/s when reaching the ATN+/AD and glued to the skull. Optic fibers were lowered vertically or in diagonal above the PreS at similar rates. For the multi-wire electrodes, the ground silver wire was implanted at the surface of the lateral cerebellum. For silicone probes, the external reference and ground wires were twisted together and implanted at the surface of the lateral cerebellum. Carprofen (5 mg/kg, i.p.) and paracetamol (2 mg/mL, drinking water) were provided during the peri-operative period. The mice were left in their home cage for a week to recover from the surgery and their weight, behavior and all aspects were monitored in score sheets established with the veterinary protocols. During this period, mice were also habituated to the handling and the recording cables.

#### Unit recordings and HD monitoring

Mice were placed into a large cylindrical Plexiglas cage (diameter: 50 cm, height: 40 cm) where they could freely behave all along the recording sessions. The cage was positioned below a vision color camera inside a Faraday cage. Implanted animals were connected to the pre-amplifier PZ5-32 (Tucker-Davis Technologies (TDT)) via a ZIF-Clip Headstage adaptor (TDT, ZCA-OMN16) for the multi-wire electrodes and a ZIF-Clip Headstage (TDT) for the silicone probes. The camera was connected to a RV2 collection device (TDT) capable of tracking red and green LEDs mounted on the ZIF-Clip Headstage. The preamplifier was connected to a main amplifier RZ5D (TDT). The main computer (WS8, TDT) used the Real-time Processor Visual Design Studio (RPvdsEx) tool to design the recording sessions, activate light stimulation from the PlexBright Optogenetic Stimulation System (Plexon), and acquire the electrophysiological data from the headstage and tracking data from the camera.

For optogenetic stimulation of the PreS/RSC (in 3 multiwire-implanted and 4 silicone probe-implanted mice), a recording session consisted in a 10 – 20 min baseline recording followed by a 10 – 20 min recording with optogenetic activation of the PreS/RSC. The stimulation consisted in 300 – 600 light stimulations of 10 ms duration, one stimulation every 2 s. The intensity of the light ranged from 2 – 6 mW depending on the quality of the homemade optic fibers. For chemogenetic silencing of the anterodorsal TRN (in 5 silicone probe-implanted mice), a recording session consisted in a 10 – 20 min baseline recording, i.p. injection of CNO (1 – 2 mg/kg) or NaCl, 40 min resting in homecage and 10 – 20 min test recording. The timing of the CNO injection is based on previous *in vivo* work using the same mouse line and CNO products, showing that the CNO effect peaked ∼30 min post i.p. injection (Fernandez et al., 2018). For the combination of both optogenetic activation of PreS/RSC and chemogenetic silencing of the anterodorsal TRN (in 3 silicone probe-implanted mice), the recording sessions consisted of 10 min baseline, 10 min optogenetic stimulation, i.p. injection of CNO (1 – 2 mg/kg), 40 min resting in homecage, 10 min second baseline and 10 min optogenetic stimulation. The number of recording sessions per mouse ranged from 1-3.

#### Spike sorting

The Offline Sorter software (Plexon), Neuroexplorer (Nex Technologies) and MATLAB (MathWorks) were used to sort and analyze single-unit spikes. The waveforms were manually delineated in the two-dimensional space of principal components using their voltage features. Single units were defined as discrete clusters of waveforms in the principal component space, and did not contain spikes with a refractory period less than 1 ms. The quantification of the clusters separation was further measured with multivariate ANOVA and J3 statistics. Cross-correlation analyses were used to control that a single unit was not recorded on multiple channels. Target units that had a peak of spike discharge when the reference unit fired were considered as duplicates and only one of the copy was used for analysis (Adapted from (Rozeske et al., 2018)). To compare the recordings during baseline and after injection of CNO or NaCl, units were sorted with two different methods. At first, both recording sessions were manually sorted as described above while the experimenter was blind to the baseline and CNO/NaCl condition. In a second step, the baseline sorting template was used for the CNO/NaCl recording. Both methods gave similar results and only the manual sorted data are shown. Some units were stable across sessions over several days or weeks, as evident by their identical firing rate, preferred direction and their detection on the same recording channel. In such cases with duplicates, only the first recording session was kept. Results did not change when analysis of later sessions were used.

#### Unit analysis

The discharge pattern of well-defined single units in the ATN+ was aligned to the optogenetic stimulation using peri-event raster plots and cumulative histogram (5 ms bins, starting 50 ms before LED onset and lasting 200 ms after LED onset, Neuroexplorer). The firing rate 50 ms before the LED onset was used as a baseline to calculate the z-score of each bin as follows: 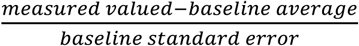. Z-scores were considered significant when > 1.96 and < -1.96. Significant changes in the firing rate fell into 4 distinct classes depending on the direction of the change (increase or decrease firing) and the timing of the change. The latencies were measured from LED onset to the first bin showing a significant z-score value. In Fig. 5K, the averaged z-score was measured from the first significant bin to the last significant bin. This range included sometimes bins that were below significance and were sparsely distributed within the evoked event. The range of bins included to quantify decrease or early increase in firing rate was identical for the baseline and the CNO recordings and based on the earliest and last significant bin regardless of the condition.

Using a custom-made Matlab routine, the discharge patterns of ATN+ units were binned to the HD of the mice. The angles of direction were binned in 6°. The firing rate was averaged for each of the 60 portions of the circle. The length of the Rayleigh vector (r) was calculated and units were considered as HD if r ≥ 0.4, as head-modulated if 0.2 ≤ r < 0.4 and as not tuned if r < 0.2 (Yoder and Taube, 2009). The maximal firing rate, the width and the preferred direction were calculated for HD units. The width of the tuning curve was measured as the span of the angle between the two directions for which the firing rate was equal to 50 % of the maximal firing rate at the preferred direction (Blair and Sharp, 1995).

### Behavioral experiment

#### Recording

One week before the beginning of the behavioral task, VGAT-Ires-Cre mice expressing either an hM4D or a non-hM4D -related (control mice) construct into the anterodorsal TRN were habituated to the handling and i.p. injection. Naïve VGAT-Ires-Cre male mice were trained to find a 12 cm wide circular platform submerged 0.5 – 0.8 cm below the surface in a 150 cm diameter circular pool filled with white opaque water at 23±1°C. Mice were trained in daily sessions composed of 4 consecutive trials, with a 60 s probe session without the platform preceding the first trial session every odd days. A trial ended when the mice spent 5 s onto the platform. Mice were left 10 s more before being placed below a heating lamp before the next trial, 10 – 15 s later. Four shapes around the pool (cross, horizontal stripes, vertical stripes, coffee grain) served as visual cues and were placed in the SW, NW, NE, SE corner of the room respectively. If the mouse failed to find the platform after 60 s, the experimenter guided it to the platform where it was left for 15 s. Mice were placed in the pool facing the wall. The position of pool entry was randomly shuffled every day between NE, SE, NW and NE. During a 60 s probe session, the platform was removed and mice were released from the wall of the quadrant opposite to the target one. The experimenter was blind to the condition of the mice (control or hM4D). The session duration (between the first and the last animal) was ∼2 hours, the first trial starting at Zeitgeber time 0 + 1.5 h. Daily i.p. injection of CNO (1 – 2 mg/kg) were performed 40 min before the beginning of the session. The timing of the CNO injection is based on previous *in vivo* work using the same mouse line and CNO products, showing that the CNO effect peaked ∼30 min post i.p. injection (Fernandez et al., 2018).

#### Analysis and automatic strategy detection

The video tracking data were analyzed using EthoVisionXT14 (Noldus) to quantify the average swimming speed, escape latency, proximity (mean distance of all the tracked points of the path to the platform center), percentage time spent in target quadrant and platform crossings. Heatmaps were generated by superimposing all the path points of every mouse in a group. Heatmaps were linearly scaled using the global minimum and maximum for both groups to allow comparison between the two. To attribute a specific strategy to each MWM trial, we used a homemade matlab algorithm based on (Garthe et al., 2009). For each trial, the animal path in the MWM was extracted as timed-tagged x and y coordinates from which specific variables were computed in order to take a decision. The 8 strategies are described in Suppl. Fig. 5 and the decision was made in the following sequential order with the 4 allocentric strategies first followed by the 4 egocentric strategies: 1-Direct swimming; if 95 % of the time-points are spent in the goal cone (isosceles triangle with its height going from starting point to goal platform with an origin angle of 40°). 2-Focal search; if the mean distance of the path to its centroid (MDTC) was inferior to 35 % standard unit (STDU) corresponding to the radius of the MWM, and the mean distance to the edge of the goal platform was inferior to 30% STDU. 3-directed search; if total time spent in the goal cone was superior to 80 %. 4-perseverance; if the MDTC was inferior to 45 % STDU and the mean distance to the previous platform edge was inferior to 40 % STDU. In our case, the perseverance strategy was only possible after day 10, during the reversal learning period. 5-chaining; if the time spent in the annulus zone (spanning from 33 to 70 % STDU) was superior to 80 %. 6-scanning; if the total coverage of the MWM (the pool was divided in 15 cm squares and the coverage was obtained as the ratio of crossed squares over the total number of squares) was superior to 10% and inferior to 60 %, and the mean distance of the path to the center of the MWM was inferior to 70% STDU. 7-thigmotaxis; if the time spent in the closer wall zone (spanning from 87 % STDU to the edge of the MWM) was superior to 35 % and the time spend in the wider wall zone (spanning from 70 % STDU to the edge of the MWM) was superior to 65 %. 8-random search; if the total coverage of the MWM was superior to 60 %. If none the conditions could be met in this order, no strategy were attributed.

#### Histology

At the end of the recording sessions, mice were perfused as described in the previous section: Perfusion and tissue processing. Brain sections were directly mounted on slides and observed though a fluorescent stereomicroscope (Nikon SMZ 25) and NIS-Elements 4.5 (Nikon) was used to analyse images. Mice showing expression of the hM4D construct outside of the anterior sector of the TRN or only were excluded.

### Statistics

All tests were done using R programming software (2.15.0, R Core Team, The R Foundation for Statistical Computing (www.rproject.org/foundation), 2007]. The normality of the data sets was assessed using Shapiro-Wilk normality test. Comparison of two data sets were done using Student’s *t* test and paired Student’s *t* test, for non-repeated and repeated measures respectively, or their non-parametric equivalent, Mann-Whitney U test and Wilcoxon signed rank-test. Chi-square tests were used to assess whether the swimming region of mice during probe sessions of the MWM were different from the expected frequencies and the proportion of strategies used between mouse groups. 1-way/2-way (non-)repeated measure ANOVAs followed by post hoc *t* tests were used when necessary on normally distributed data sets whereas non-normally distributed data were analyzed directly with the post hoc tests. A Bonferroni correction was applied when more than two comparisons were done on the same data set and the new alpha threshold is indicated. All statistical tests are specifically indicated in the figure legends if they are not given in the main text.

