## Supplemental figures for "A thalamic reticular circuit for head direction cell tuning and spatial navigation"

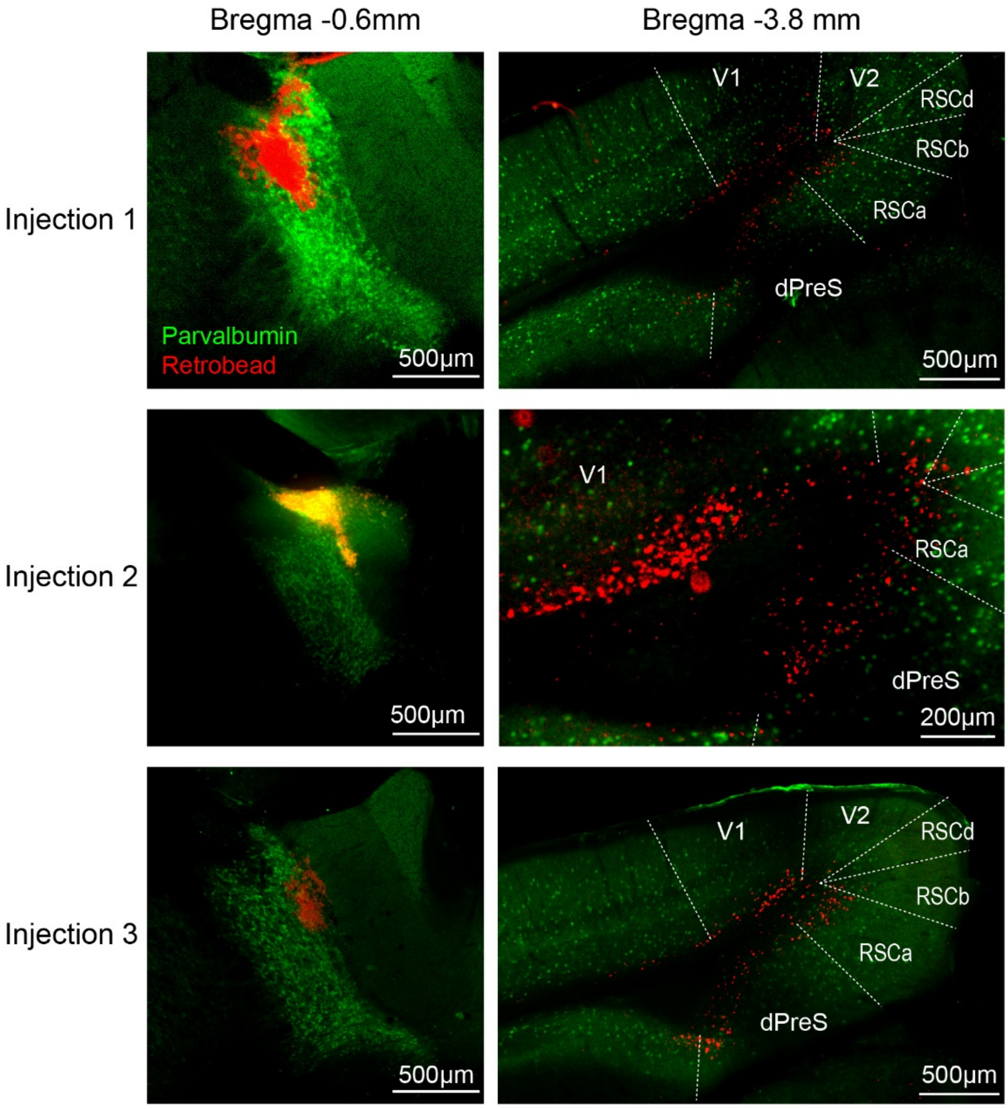


**Suppl. Figure 1. The anterior TRN receives afferents from RSC, PreS, and visual cortices.**

Left: Three epifluorescent micrographs of mouse coronal brain sections showing the site of injections of retrobeads (red) into the anterodorsal portion of the TRN. Right: Corresponding cortical structures where beads were retrogradely transported, thus indicating afferent origin. Green: immunostaining for parvalbumin. Dorsal presubiculum (dPreS) – Retrosplenial cortex (RSC) – Visual cortex (V1/V2).

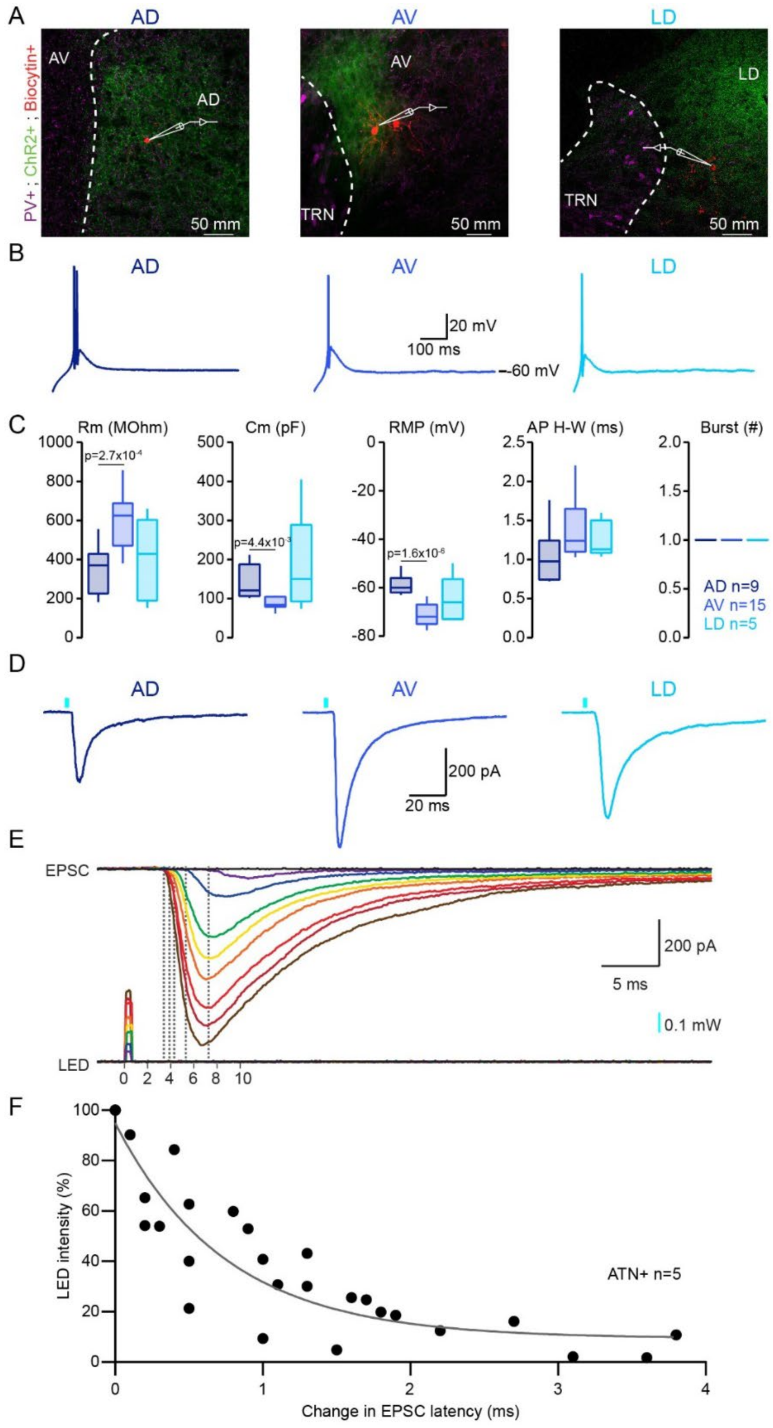


**Suppl. Figure 2. The PreS/RSC establish functional excitatory synapses onto AD, AV and LD.**

**(A)** Confocal micrographs of 300 µm-thick mouse brain sections showing the whole-cell recorded AD (left), AV (middle) and LD (right) neurons filled with neurobiotin (red). Green, ChR2-EYFP-expressing PreS/RSC afferents, magenta, PV+ TRN cells. **(B)** Voltage responses of neurons shown in A to a negative current injection. **(C)** Box-and-whisker plots of passive and active cellular properties of PreS/RSC-connected AD (green, n=9), AV (orange, n=15) and LD (blue, n=5) neurons. From left to right: Membrane resistance (Rm), membrane capacitance (Cm), resting membrane potential (RMP), action potential (AP) half-width (H-W), burst number. Student's *t* tests were used to compare Vm, Rm and Cm. A Mann-Whitney U test was used for comparing HW. **(D)** Current responses of AD (left), AV (middle) and LD (right) neurons to optogenetic activation (blue bars, 1 ms, 3.5 mW power, 455 nm) of PreS/RSC afferents, recorded at -60 mV. **(E)** Current responses of a thalamic neuron to optogenetic stimulation of PreS/RSC afferents at different intensities. **(F)** Graph showing the increasing latency of the EPSC latency when reducing light stimulation intensity in 5 ATN+ neurons.

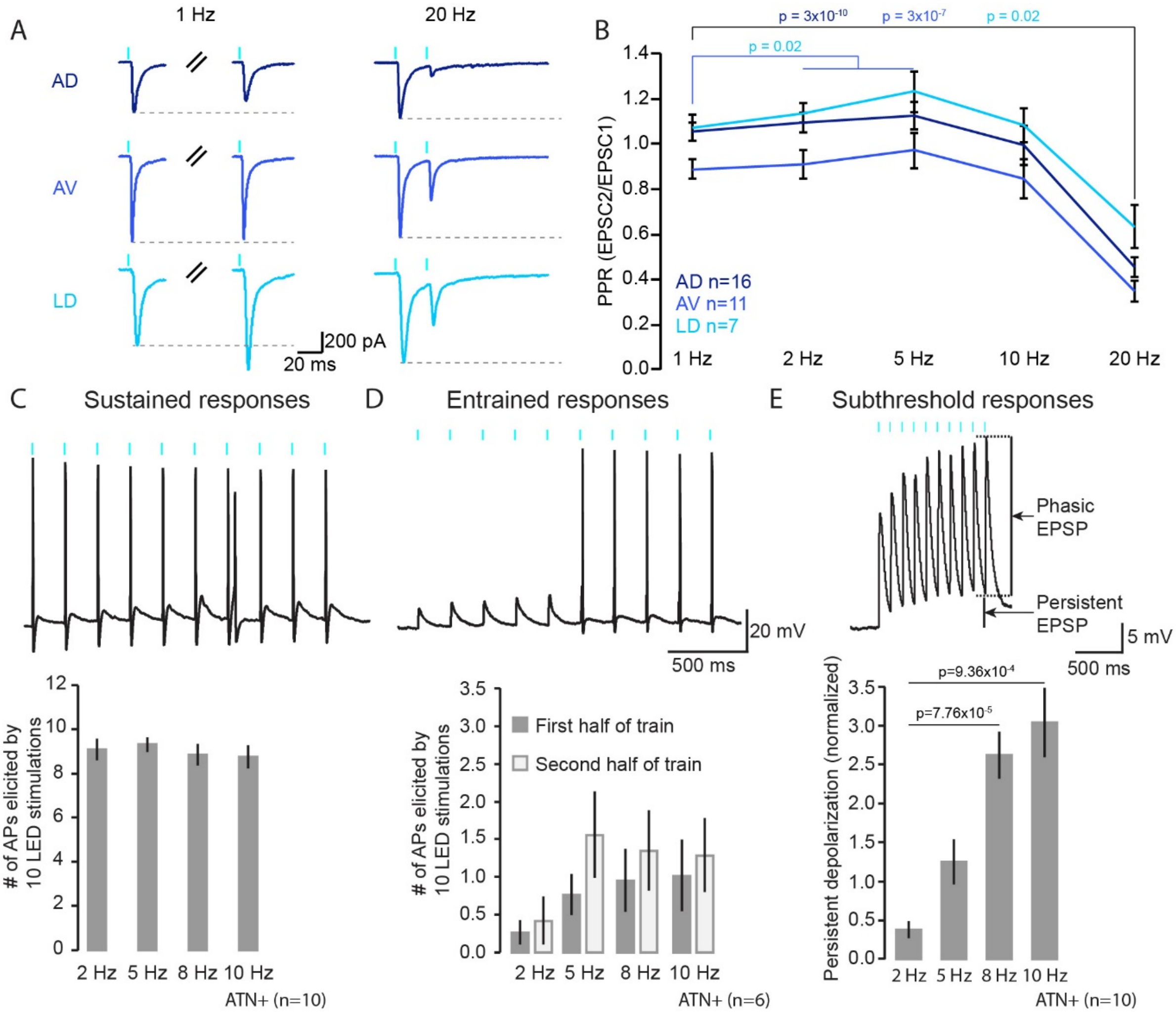


**Suppl. Figure 3. The ATN+ receives strong unitary connections with 'driver' characteristics from the PreS/RSC.**

**(A)** Representative current responses of AD, AV and LD neurons held at -60 mV upon paired-pulse stimulation at 1 and 20 Hz. Grey dotted lines correspond to the amplitude of the first EPSC. **(B)** Graph of the paired-pulse ratios. Paired Student's *t* tests with Bonferroni correction for multiple comparisons for PPR at 1 Hz vs PPR at other frequencies ($\alpha = 0.013$). **(C)** Top: typical membrane voltage response of an ATN+ neuron to a train of 10 light stimulations at 10 Hz. Action potentials are elicited from the first light stimulation, classifying it as a sustained response. Bottom: Histogram of action potential numbers during such train stimulations in ATN+ neurons (n = 10) showing sustained response patterns. Wilcoxon signed rank-tests were used to compare between the 2 Hz and the other frequencies of stimulation, with Bonferroni correction for multiple comparison ($\alpha = 0.017$). **(D)** Top: same as in C when neurons responded with a subthreshold response at train onset, classifying them as an entrained response. Bottom: Histogram of action potential numbers during the first half (5 first light stimulation) and the second half of the train of 10 light stimulations in ATN+ neurons (n = 6). A Wilcoxon signed rank-test (at 2 Hz) and Paired Student's *t* tests (at 5, 8, 10 Hz) were used to compare the number of action potentials elicited during the first half vs the second half of the train of stimulation. **(E)** Top: same as in C for subthreshold responses in an ATN+ neuron held at -60 mV. Bottom: Histogram of the persistent depolarization induced by trains in TRN neurons (n = 5). The persistent depolarization was measured on the last 3 stimulations of the train, see methods for the details. Paired Student's *t* tests with Bonferroni correction for multiple comparison ($\alpha = 0.017$) were used to compare the persistent depolarization at 2 Hz with the persistent depolarization at other frequencies.

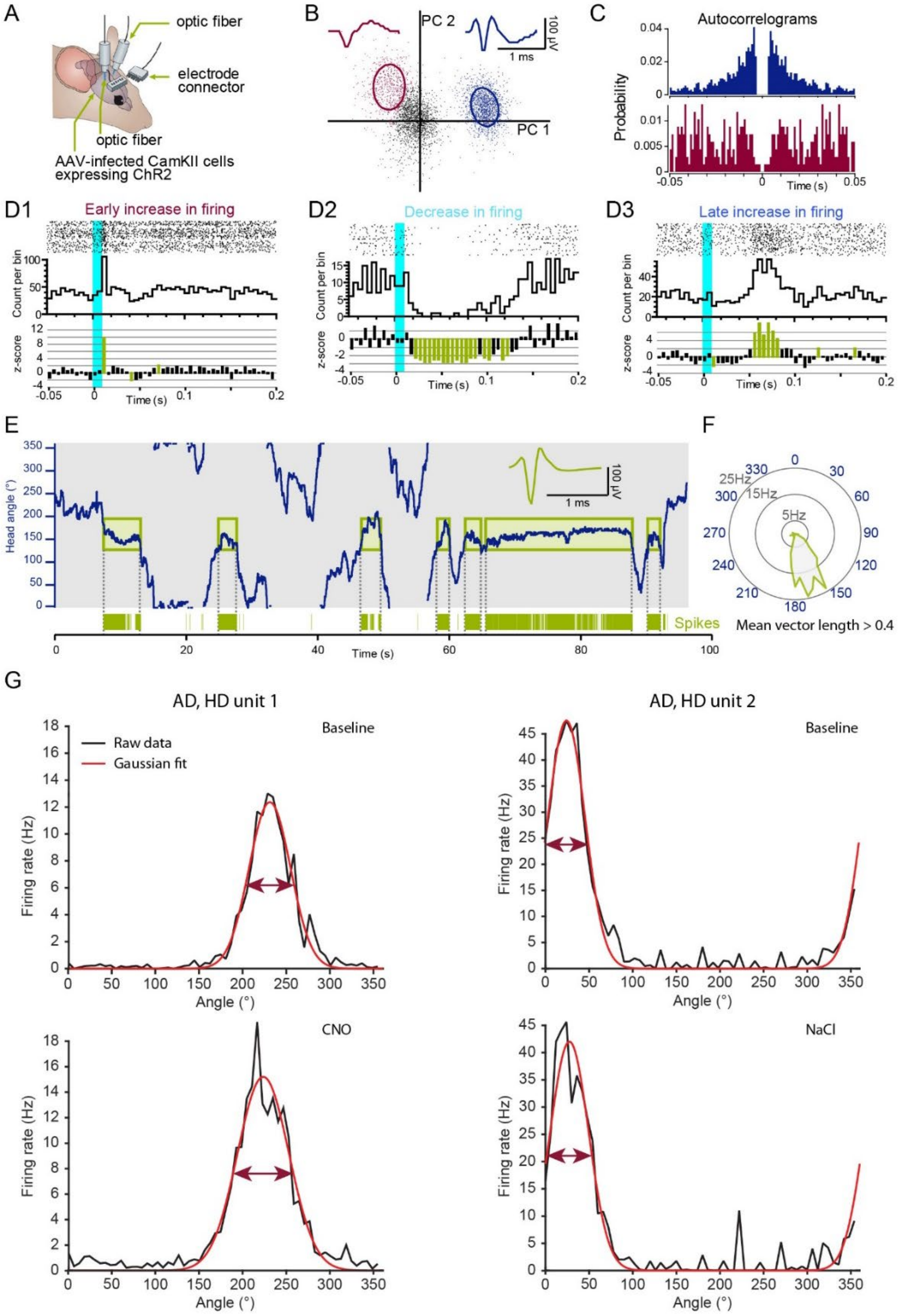


**Suppl. Figure 4. Sorting and analysis of the *in vivo* single unit recording.**

**(A)** Scheme of the *in vivo* freely moving recording configuration. **(B, C)** Example of unit sorting based on principal component analysis (B) and autocorrelation showing the typical refractory period around 0 (C). **(D1-D3)** Raster plot, cumulative histogram and z-score analysis for three characteristic unit responses. **(E)** Graph showing the mouse HD (blue trace) in combination with detected spikes of a putative thalamic HD unit (vertical green lines) tuned around 150°. Insets: mean unit waveform. **(F)** Polar plot of the tuning curve of the HD unit shown in (E). Units with a mean vector length above 0.4 were considered as HD units. **(G)** Graphs of the raw firing rate of two AD HD units (black) and the corresponding Gaussian fit (red) shown in figure 5G. Top: baseline firing rate. Bottom left: firing rate 40 min after CNO i.p. injection. Bottom right: firing rate 40 min after NaCl i.p. injection.

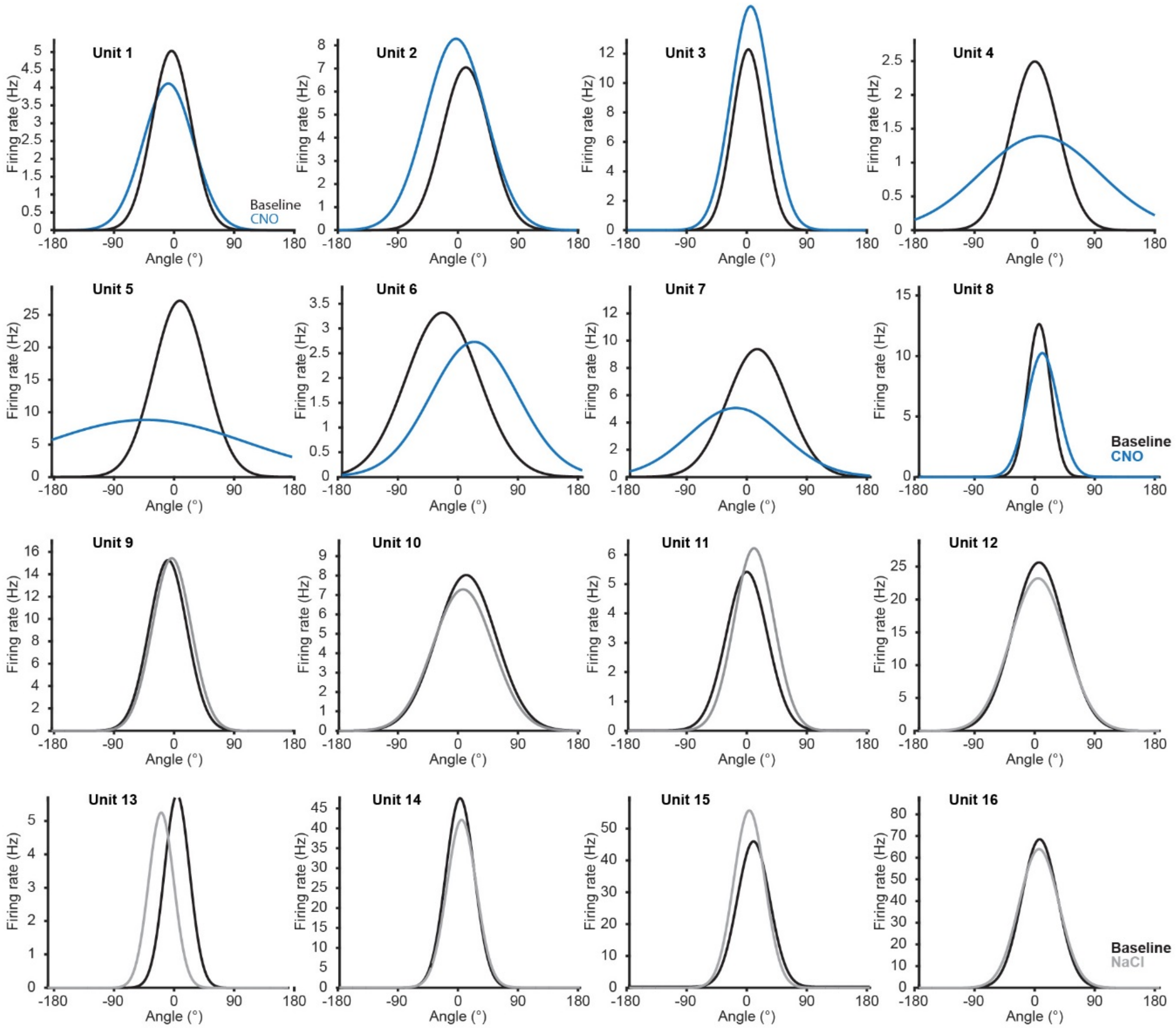


**Suppl. Figure 5. Effect of chemogenetic silencing of anterodorsal TRN neurons onto the tuning of AD HD units.**

Graphs of the discharge pattern of 16 AD HD units from five VGAT-Ires-Cre mice expressing the hM4D in the anterodorsal portion of the TRN. The line shows the Gaussian fit of the firing rate of the HD units during baseline recording (black). 40 min after CNO i.p. injection (blue) or 40 min after NaCl i.p. injection (grey).

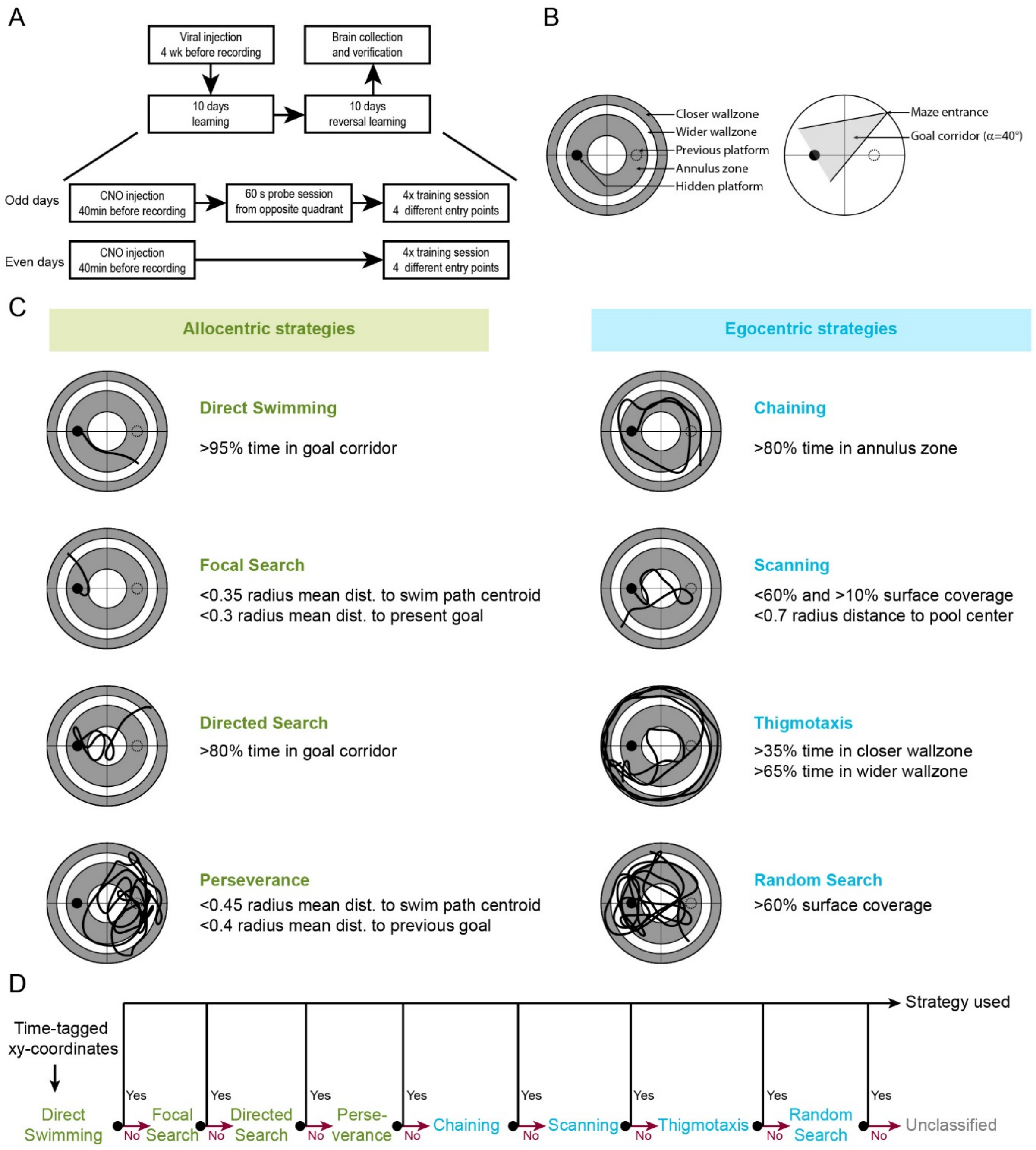


**Suppl. Figure 6. Design of Morris Water tasks and algorithm-based classification of search strategies.**

**(A)** Scheme of the experimental design. For further details, see methods. **(B)** Representation of the variables used for the classification process. The pool was divided into several areas to calculate the respective amount of time spent into these specifics areas (left). The average distance of all the data points constituting the search path to its centroid, the present platform and the former platform was used for the classification. Search patterns based on a directional preference for the actual platform were identified using a triangular shaped corridor expanding from the entry point of the mice with its bisecting line towards the platform (right). **(C)** Strategies were identified by one or two parameters representing their major properties. **(D)** The algorithm excluded strategies from the more to the less specific search patterns. Search patterns not recognized were grouped as unclassified strategies.

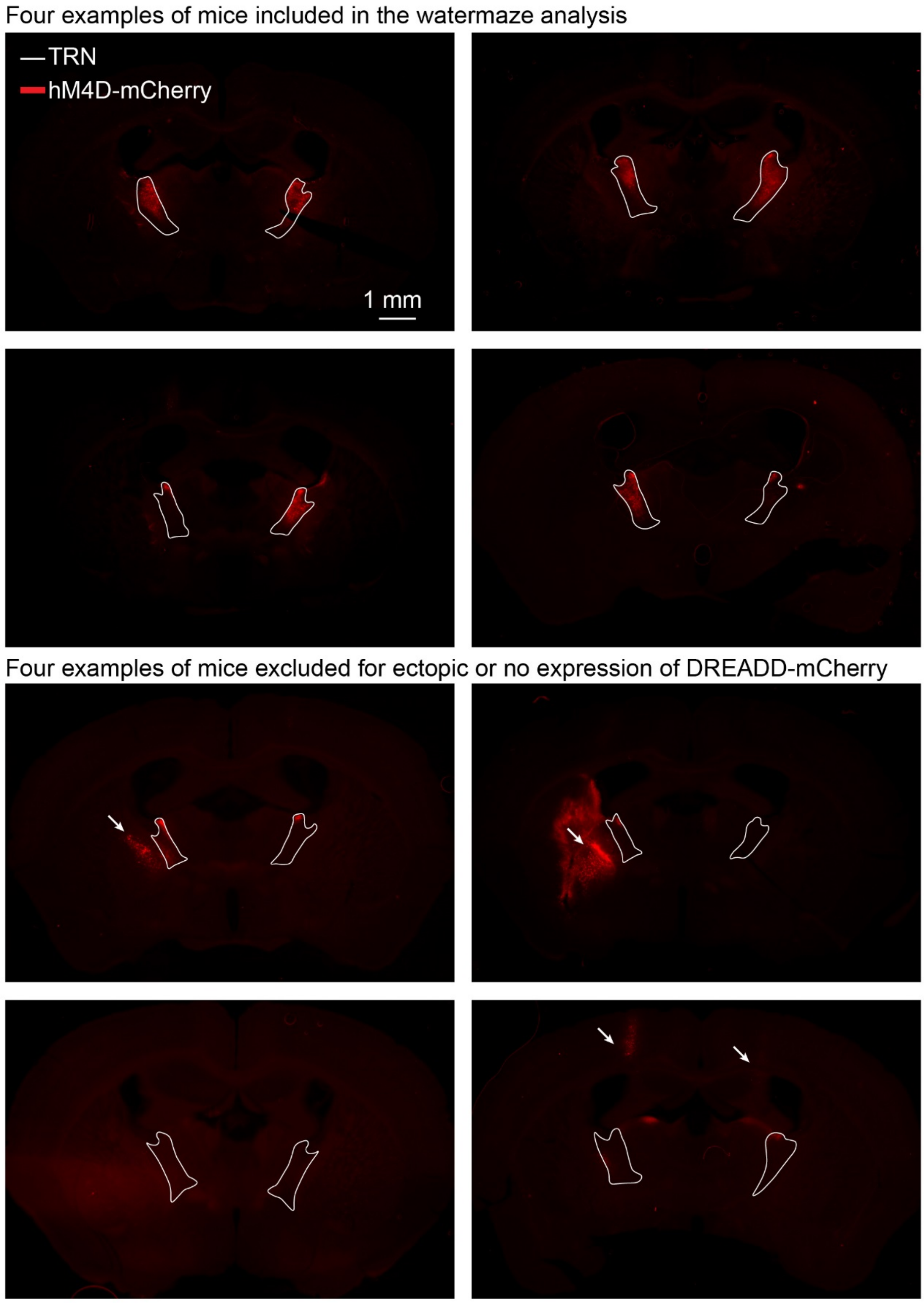


**Suppl. Figure 7. Expression of hM4D in the anterodorsal TRN of VGAT-Ires-Cre mice trained in the Morris Watermaze.**

Eight epifluorescent micrographs showing the expression of hM4D in VGAT-Ires-Cre mice. The top four mice were included in the analysis as the hM4D is expressed specifically in the anterodorsal portion of the TRN. The bottom four mice were excluded from the analysis because hM4D was expressed in the Globus Pallidus (top left and right), not expressed in the anterodorsal TRN (bottom left) or expressed in the cortex (bottom right).
